# Sustained darkness reveals rapid phenotypic accommodation and loss of plasticity in cavefish evolution

**DOI:** 10.64898/2026.04.28.721295

**Authors:** Laura Guerrero Pena, Marina Horvatiček, Mateo Čupić, Marko Lukić, Tin Rožman, Helena Bilandžija

## Abstract

How environmentally induced traits become genetically stabilized during evolution remains a central question in biology. The teleost *Astyanax mexicanus*, comprising surface (SF) and cave (CF) morphotypes, provides a powerful system to link environmentally induced traits with evolved phenotypes. Here, we integrate morphometric, physiological, and transcriptomic analyses to investigate how sustained darkness reshapes phenotypic plasticity across generations.

We show that plastic responses in SF are rapidly modified after only two generations of dark rearing (2dSF). Morphological changes in 2dSF are highly heterogeneous: some traits shift toward cavefish values, while others overshoot them, and key traits such as eye size and pigmentation respond in the opposite direction, indicating widespread maladaptive plasticity. In contrast, CF exhibit limited environmental responsiveness, consistent with canalized phenotypes. Traits exhibiting maladaptive plasticity in SF tend to be canalized in CF, suggesting that maladaptive responses are preferentially reduced during evolution.

At the molecular level, transcriptional plasticity declines sharply from SF to 2dSF and is nearly absent in CF, suggesting rapid loss of environmental responsiveness. Gene expression analyses further reveal a mixture of maladaptive and adaptive trajectories. Most transcriptional changes in 2dSF deviate from cavefish expression patterns, including those related to visual function, consistent with maladaptive morphological trends. However, a subset of genes involved in lipid metabolism and heme homeostasis shows stepwise shifts from SF through 2dSF to CF, consistent with adaptive accommodation. Notably, anesthesia resistance increases in 2dSF to levels comparable to CF within only two generations, demonstrating rapid accommodation in neural traits.

Together, our results show that plastic phenotypes are initially heterogeneous but are rapidly reshaped across generations, with maladaptive components reduced and adaptive components stabilized. This process bridges immediate plasticity and long-term evolutionary change, providing empirical support for phenotypic accommodation as a mechanism facilitating rapid adaptation and highlighting how plasticity can biasing evolutionary trajectories during the colonization of extreme environments.

## INTRODUCTION

Phenotypic plasticity, the ability of organisms to alter their phenotype in response to environmental conditions, is a ubiquitous property of living systems (1,2). Because plastic responses can arise immediately, they are often proposed as a central feature of how populations cope with environmental change. However, whether plasticity merely buffers organisms against environmental change or actively contributes to evolutionary divergence and genetic adaptation remains a matter of debate (3–5). A key unresolved issue is how an initially plastic response could be transformed into a stable, evolved phenotype, and the empirical data from the early stages following environmental transition is needed (6).

The plasticity-first (6) or genes as followers hypothesis (7) proposes that phenotypic plasticity contributes to evolutionary change by generating immediate plastic responses, followed by subsequent modification of those responses, and the emergence of canalized phenotypes that are insensitive to environmental variation. Natural selection can act on phenotypic variation revealed by environmental shifts, refining and stabilizing the phenotype over subsequent generations, and this process is termed phenotypic and genetic accommodation. The final stage is genetic assimilation, where traits become constitutively expressed and no longer depend on the original environmental cue (1,4,8–11). Although these processes are often discussed conceptually, empirical studies that disentangle them especially within the same species, and on short evolutionary timescales, remain scarce. In particular, it is largely unknown how rapidly plastic responses themselves can change across generations, and whether canalization represents an environmentally inducible state or an evolutionary endpoint. Addressing these questions is essential for evaluating the generality of plasticity-first models across diverse systems experiencing persistent environmental change.

The most compelling evidence of plasticity-first evolution would be to observe it taking place in organisms where ancestral conditions are known, environmental drivers are identified and the environmental change occurs rapidly (6). Cave colonization provides a rare natural setting in which all these criteria are fulfilled. The transition from surface to the cave represents an extreme environmental shift: caves are characterized by perpetual darkness, food scarcity, and altered biotic interactions. These strong selection pressures result in a consistent suite of traits that evolved independently in cave-dwelling species across diverse phyla, from planaria to vertebrates (12). Among the diverse cave-dwelling animals, the Mexican tetra (*Astyanax mexicanus*) stands out as a unique model system since it includes both the ancestral surface-dwelling (SF) and multiple independently evolved cave-dwelling (CF) populations. In a short evolutionary time of only 30,000-200,000 generations (13,14), cavefish evolved striking phenotypic alterations, including the loss of eyes and pigmentation, enhanced non-visual sensory systems, altered metabolism, and nervous system changes, raising the question of mechanisms that enable rapid evolution of such a diverse set of traits.

In this study, we present new experimental evidence that bridges the gap between environmentally induced plasticity and genetically assimilated adaptation. By rearing *A. mexicanus* surface fish under sustained darkness across successive generations (Fig 1), we observe that traits become modified beyond their initial plastic responses, providing empirical support for phenotypic accommodation. Furthermore, cavefish demonstrate robust, mostly environmentally insensitive expression of the same traits, consistent with genetic assimilation. Our results suggest that plasticity-led evolution can proceed over remarkably short timescales even in laboratory conditions and offer a possible mechanistic basis for the repeated, rapid emergence of cave-adapted phenotypes in *A. mexicanus*.

**Fig 1.**
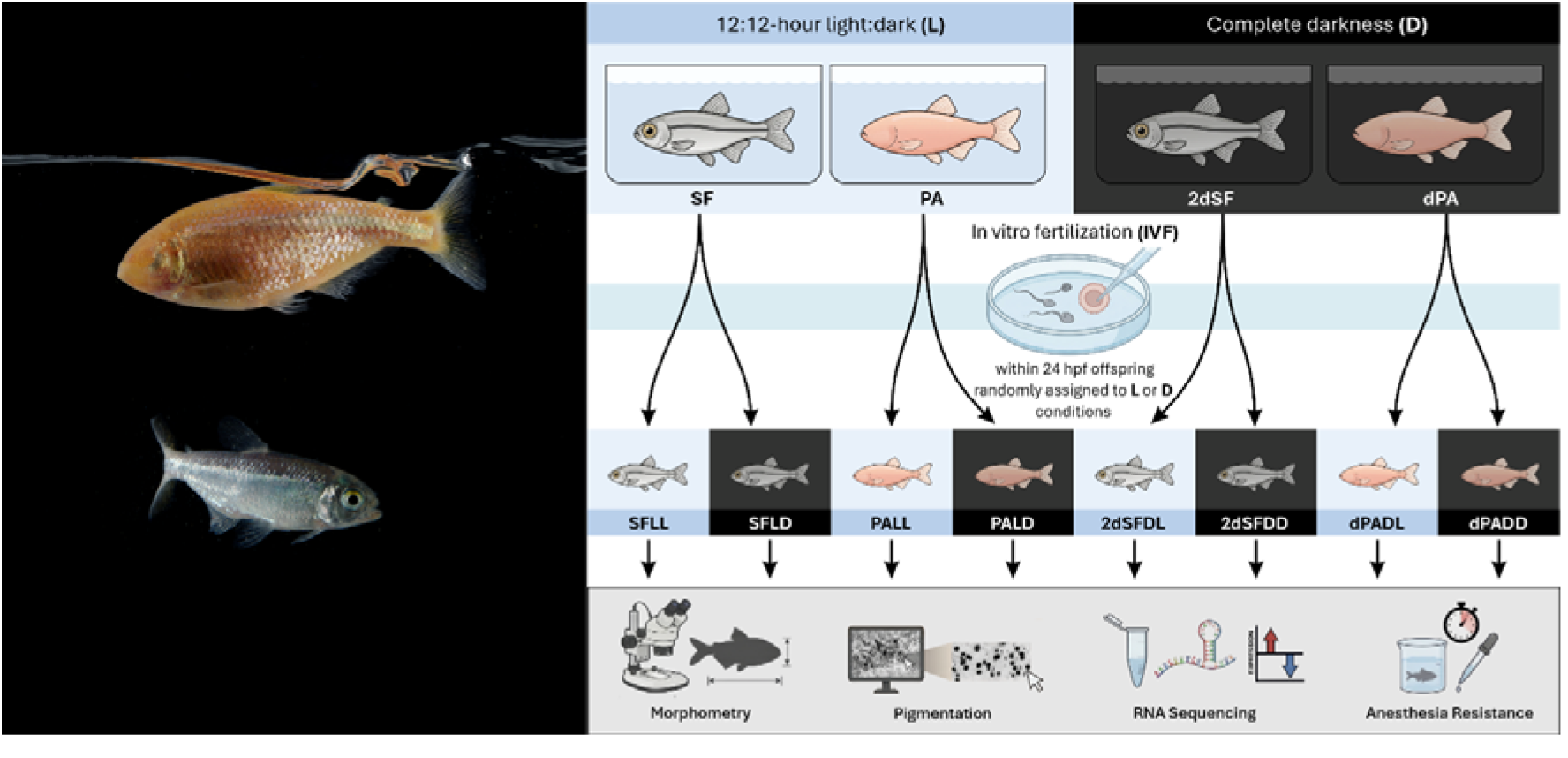
Study system and experimental design. (A) Two morphotypes of the Mexican cave tetra *Astyanax mexicanus*: surface fish with well-developed eyes and pigmentation, and Pachón cavefish lacking eyes and pigmentation. (B) Schematic representation of the experimental design.

## RESULTS

### Surface and cave morphotypes remain distinct across light environments

A discriminant function analysis (DFA) conducted on 239 *A. mexicanus* larvae across eight experimental groups (N = 30 for each group except dPADL; N = 29) identified seven discriminant functions. SF and PA are completely separated by the first function (Fig 2A), which explains 99.4% of variance (eigenvalue=166.80, r=0.997). Significance testing shows that functions 1-4 are statistically significant (p<0.003). The structure matrix shows that the first function is defined by eye morphology: APED (r=−0.548) and DVED (r=−0.546) and the second function by BHH (r=0.692), FINS (r=0.579), and JAHS (r=0.561) (S4 Fig). Overall classification accuracy is 61.9% for original classification and 49.8% for cross-validation. Importantly, discrimination between PA and SF populations was 100% accurate (S1 Table).

**Fig 2.**
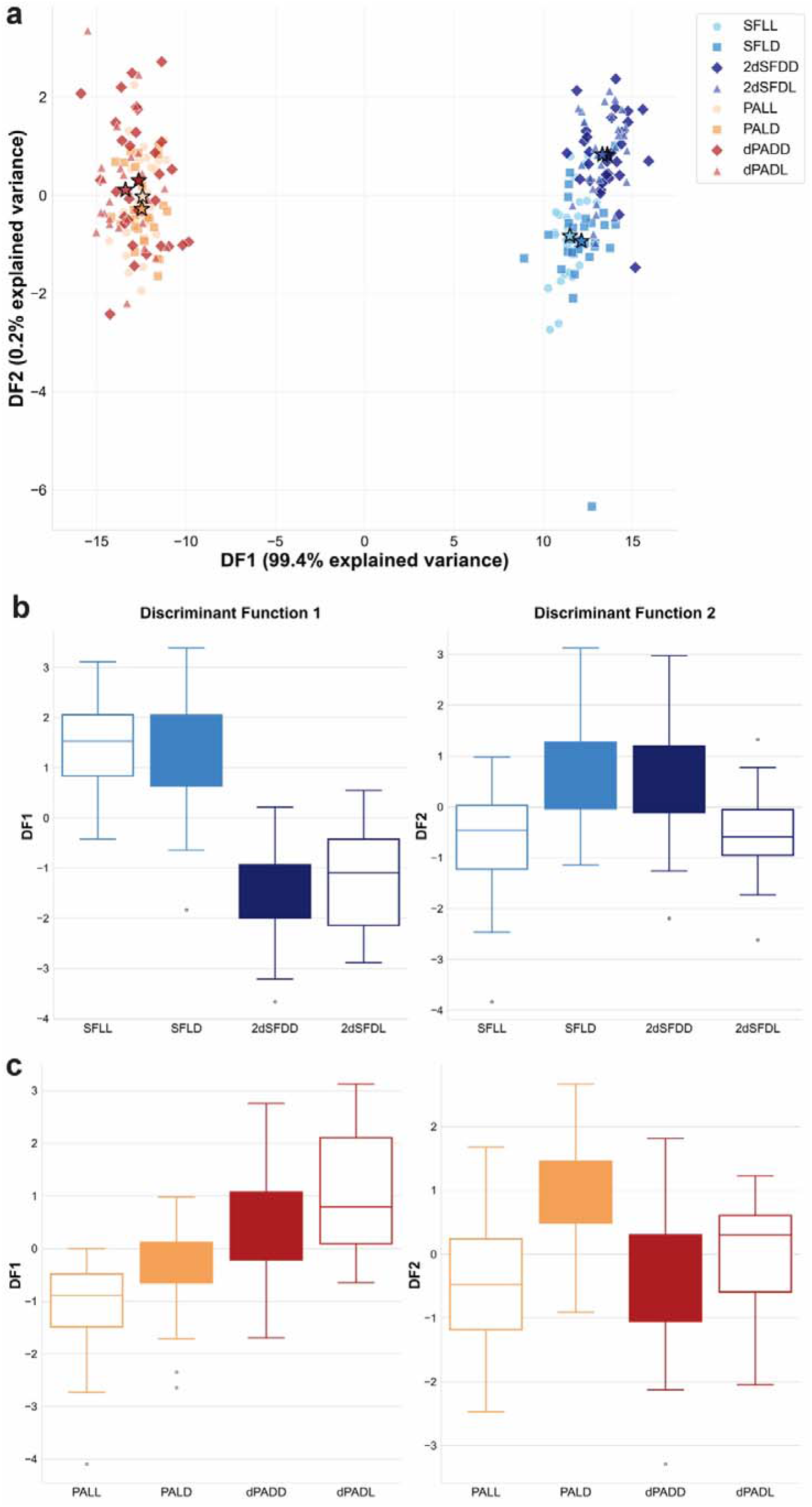
Morphometric discrimination of 7 dpf *A. mexicanus* surface fish (SF) and Pachón cavefish (PA) across light regimes. Fish were raised under four light regimes: LL, light–dark cycle; DD, constant darkness; LD, switched from light–dark to darkness; and DL, switched from darkness to light–dark, with switches applied within 24 hpf. dPA denotes PA reared in darkness for one generation and 2dSF denotes SF reared in darkness for two generations. N = 30 for all groups except PA-DL (N = 29). (A) Discriminant function analysis (DFA) scatterplot showing separation of SF and PA along discriminant functions 1 and 2. (B) Boxplots of the first two discriminant functions for SF. (C) Boxplots of the first two discriminant functions for PA.

### Plasticity differs between surface and cave morphotypes

To understand the effect of environment on *A. mexicanus* SF and PA which might otherwise be missed in a full dataset, we conducted DFA for each morphotype separately. DFA on 120 SF larvae across four experimental light conditions identified three discriminant functions. The first function explained 81.4% of variance (eigenvalue=1.86, r=0.806), the second function explained 15.2% of variance (eigenvalue=0.35, r=0.507), and the third function explained 3.4% of variance (eigenvalue=0.08, r=0.268). Functions 1-3 and 2-3 were statistically significant (1-3: Wilks’ Λ=0.241, χ^2^(36)=157.91, p<0.001; 2-3: Wilks’ Λ=0.689, χ^2^(22)=41.32, p=0.008) (Fig 2B). The structure matrix shows that the first function is most strongly defined by eye size: DVED (r=0.707) and APED (r=0.659), and traits related to head morphology JAHS (r=0.345), and HL (r=0.301) and body height BHH (r=0.500) (S5 Fig). This function clearly reflects the influence of long-term rearing conditions. The second function is most strongly defined by FINS (r=0.419), DEYE (r=0.386), and BW (r=0.269) and reflects short-term rearing environments. Overall classification accuracy was 70.8% for original classification and 52.5% for cross-validation. Correct classification for different groups ranged from 76.7 to 66.7%. The misclassification pattern shows that individuals were predominantly assigned to groups sharing the same long-term rearing condition (e.g., 2dSF-DD→2dSF-DL: 33.3%, SF-LL→SF-LD: 20.0%), indicating that parental light environment has lasting effects on morphology (S2 Table). Seven variables show significant differences among SF groups: DVED (F(3,116)=37.16, p<0.001), APED (F(3,116)=31.28, p<0.001), BHH (F(3,116)=18.05, p<0.001), FINS (F(3,116)=12.11, p<0.001), SNM (F(3,116)=9.72, p<0.001), JAHS (F(3,116)=8.65, p<0.001), and HL (F(3,116)=7.09, p<0.001) (Fig 3A).

**Fig 3.**
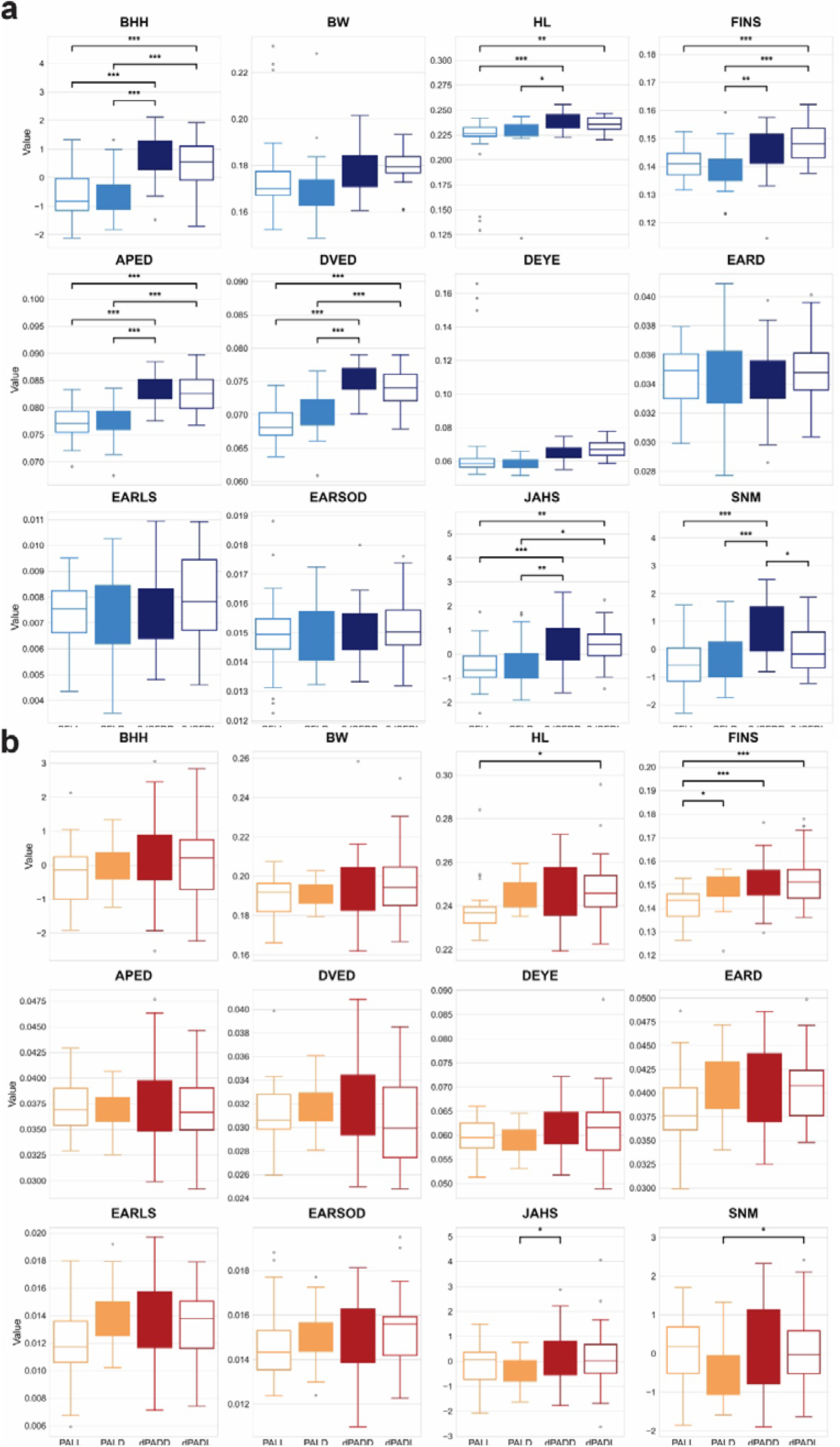
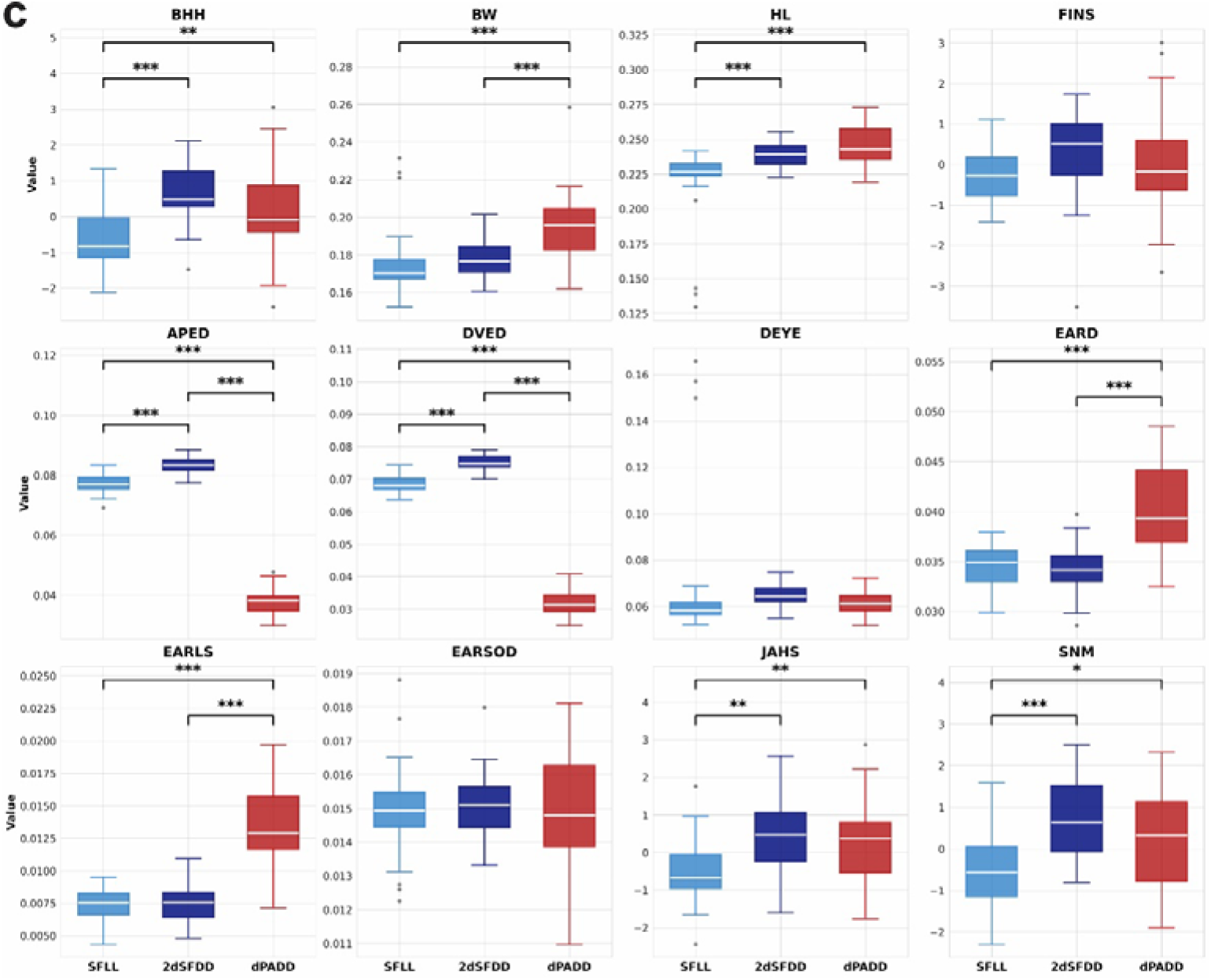
Morphometric responses of 7 dpf *A. mexicanus* surface fish and Pachón cavefish across different light regimes. (A) Surface fish (SF). (B) Pachón cavefish (PA). Fish were raised under the same light regimes described in Fig 2. Environmental switches were applied within 24 hpf. dPA indicates PA reared in darkness for one generation and 2dSF indicates SF reared in darkness for two generations. Sample sizes were N = 30 for all groups except PA-DL (N = 29). (C) Morphometric differences between morphotypes reared in original or long-term environmental conditions: SFLL reared in standard light/dark conditions, and 2dSF and dPA in constant darkness.

DFA on 119 PA cavefish larvae across four experimental light treatments extracted three discriminant functions. The first function explained 58.9% of variance (eigenvalue=0.63, r=0.623), the second function explained 30.0% of variance (eigenvalue=0.32, r=0.494), and the third function explained 11.1% of variance (eigenvalue=0.12, r=0.326). Functions 1-3 and 2-3 were statistically significant (1-3: Wilks’ Λ=0.414, χ^2^(36)=97.11, p<0.001; 2-3: Wilks’ Λ=0.676, χ^2^(22)=43.14, p=0.005) (Fig 2C). The structure matrix shows that the first function is defined by variables related to overall body shape and fin length: FINS (r=0.592), HL (r=0.342), DEYE(r=0.250), BW (r=0.220), and BHH (r=0.186). The second function is most strongly defined by variables related to the head including jaws, snout and vestibular system: SNM (r=0.446), JAHS (r=0.299), EARLS (r=−0.293) and EARD (r=−0.288) (S6 Fig). Overall classification accuracy was 61.3% for original classification and 43.7% for cross-validation. Correct classification ranged from 76.7 to 43.3%. PA long term reared in light/dark conditions (LL and LD) showed classification accuracy similar to the SF groups (76.7 and 73.3% respectively), which was markedly better than dPA (dPA-DD: 43.3% and PA-DL: 51.7%). Moreover, misclassification patterns are not as clear as in SF. In some groups, PA individuals from the same long-term rearing environment are not assigned to their respective groups transferred to the opposite light conditions (e.g. the highest misclassification for dPADL was to PALD, and for PALD to dPADD) (S2 Table). Four variables show significant differences among PA groups: FINS (F(3,115)=9.39, p<0.001), HL (F(3,115)=3.52, p=0.017), SNM (F(3,115)=3.06, p=0.031), and JAHS (F(3,115)=2.77, p=0.045) (Fig 3B).

### Sustained darkness induces heterogeneous phenotypic shifts

To assess how sustained darkness modifies SF phenotypes across generations and how these shifts relate to the cave phenotype of PA, we compared SF reared in their standard light conditions (SFLL), with 2dSF and PA reared in darkness (2dSFDD and dPADD). Morphometric responses in 2dSF were highly heterogeneous in both direction and magnitude. Some traits remained closer to ancestral SF values, whereas others approached or exceeded PA values. Body and head height, as well as composite craniofacial variables (JAHS and SNM), shifted toward the cave phenotype but overshot PA values. Only head length closely matched PA in both direction and magnitude. In contrast, eye traits shifted in the opposite direction from the cave phenotype. Traits that did not differ between SF and PA (e.g., DEYE, EARSOD) showed no induction in 2dSF, and PA-specific traits related to ear morphology and body width were likewise unresponsive (Fig 3C). Overall, darkness-induced shifts in 2dSF were trait-specific, varying in direction, and magnitude in relation to the evolved cave phenotype.

### Pigmentation shows reversible and maladaptive plasticity in surface fish

Loss of pigmentation is a hallmark cave adaptation that has evolved independently in multiple cave populations of *A. mexicanus*. Exposure to complete darkness in ancestral SF induced a significant increase in melanophore coverage within seven days. Elevated pigmentation was also observed in 2dSF. However, when 2dSF were returned to light conditions, pigmentation decreased to levels below those measured in SF reared under standard light/dark cycles (Fig 4A). In contrast, cavefish exhibited minimal changes in pigment cell numbers across rearing environments (Fig 4B). Thus, darkness induces increased pigmentation in surface fish, in a direction opposite to the reduced pigment cell numbers in cavefish, and this response remains reversible across generations. In contrast, cavefish show limited responsiveness in pigment cell numbers.

**Fig 4.**
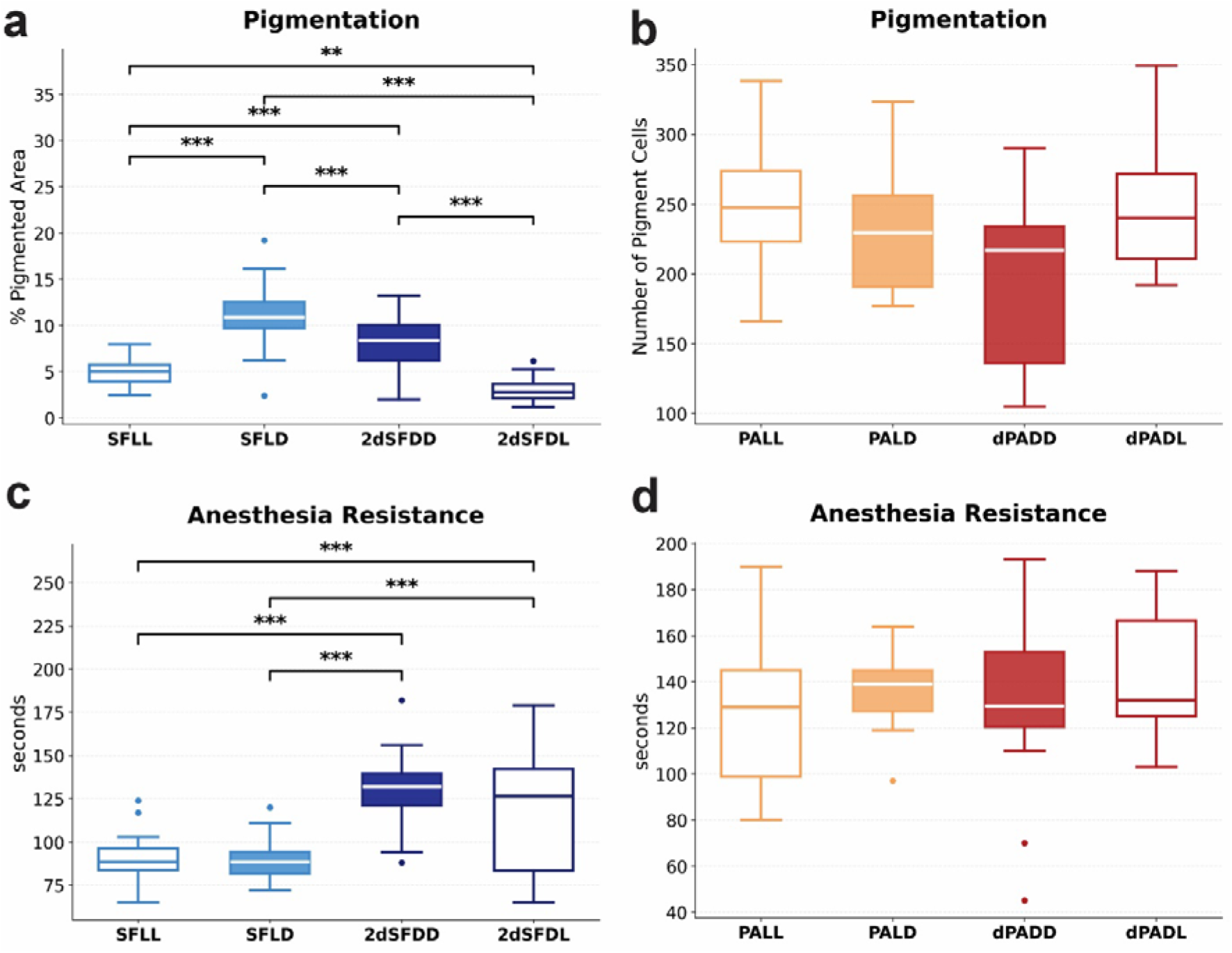
Pigmentation and anesthesia resistance in different morphotypes and rearing conditions of *Astyanax mexicanus*. Fish were raised under four light regimes: LL, light–dark cycle; DD, constant darkness; LD, switched from light–dark to darkness; and DL, switched from darkness to light–dark, with switches applied within 24 hpf. (A) Pigmentation in 7 dpf surface fish (SF) larvae, measured as percentage of pigmented area. N= 30 for all groups. 2dSF denotes SF reared in darkness for two generations. (B) Pigmentation in 7 dpf Pachón cavefish (PA) larvae, measured as number of pigment cells after L-DOPA rescue. N=10 for all groups, except dPADD where N=9. (C) Anesthesia resistance in 13 dpf larvae. N=24 for SFLL and SFLD; N=23 for 2dSFDD; N=22 for 2dSFDL. (D) Anesthesia resistance in 13 dpf PA larvae. N=24 for PALL; N=23 for PALD; N=22 for dPADD; N=7 for dPADL. Statistical comparisons by Welch’s ANOVA (A,C,D), and one-way ANOVA (B), followed by Tukey’s (A) or Games-Howell (C-D) post-hoc test.

### Anesthesia resistance increases across generations under darkness

Numerous nervous system modifications have evolved in *Astyanax* cavefish (15,16), and several cave populations exhibit elevated anesthesia resistance, a proxy for altered arousal state and neural responsiveness (17). At 13 dpf, 2dSF showed significantly higher anesthesia resistance than SF (Fig 4C), and importantly, levels were comparable to those observed in PA cavefish. In contrast, PA showed no significant differences in anesthesia resistance across treatments (Fig 4D), indicating limited environmental responsiveness. Thus, anesthesia resistance increases in surface fish under sustained darkness and converges on cavefish levels within two generations, whereas PA maintain stable levels irrespective of light conditions.

### RNA-seq data quality assessment

To investigate gene expression in *A. mexicanus*, eighteen larval transcriptomes were sequenced from three groups (SF, 2dSF and dPA) across distinct light conditions (SFLL, SFLD; 2dSFDD, 2dSFDL; dPADD, dPADL), yielding an average of 95.58 million raw reads (47.79 million read pairs) per sample. After quality filtering and adapter trimming, 97.74% of reads were retained.

High-quality reads were mapped to the AstMex3.0 reference genome (GCF_023375975.1), with an average of 29.98 million reads (64.42%) uniquely mapped per sample. Of 52,494 predicted protein-coding sequences, 97.8% were successfully annotated using eggNOG-mapper v.2.1.12, and 36,289 proteins (69.1%) were assigned at least one GO term. The functional characterization was further categorized into the three main GO domains, identifying 30,817 biological processes, 12,805 molecular functions and 4,573 cellular component terms.

### Progressive loss of light-induced transcriptional plasticity

Transcriptional plasticity was quantified by comparing short-term light versus dark responses at 7 dpf within each group (SFLL vs. SFLD, 2dSFDD vs. 2dSFDL, dPADD vs. dPADL). SF exhibited the highest environmental responsiveness, with 68 DEGs (|log2FC| > 0.5, padj < 0.05) (Fig 5A, left panel). Genes downregulated in the dark included circadian regulators, while upregulated genes were involved in metabolic and regulatory processes.

**Fig 5.**
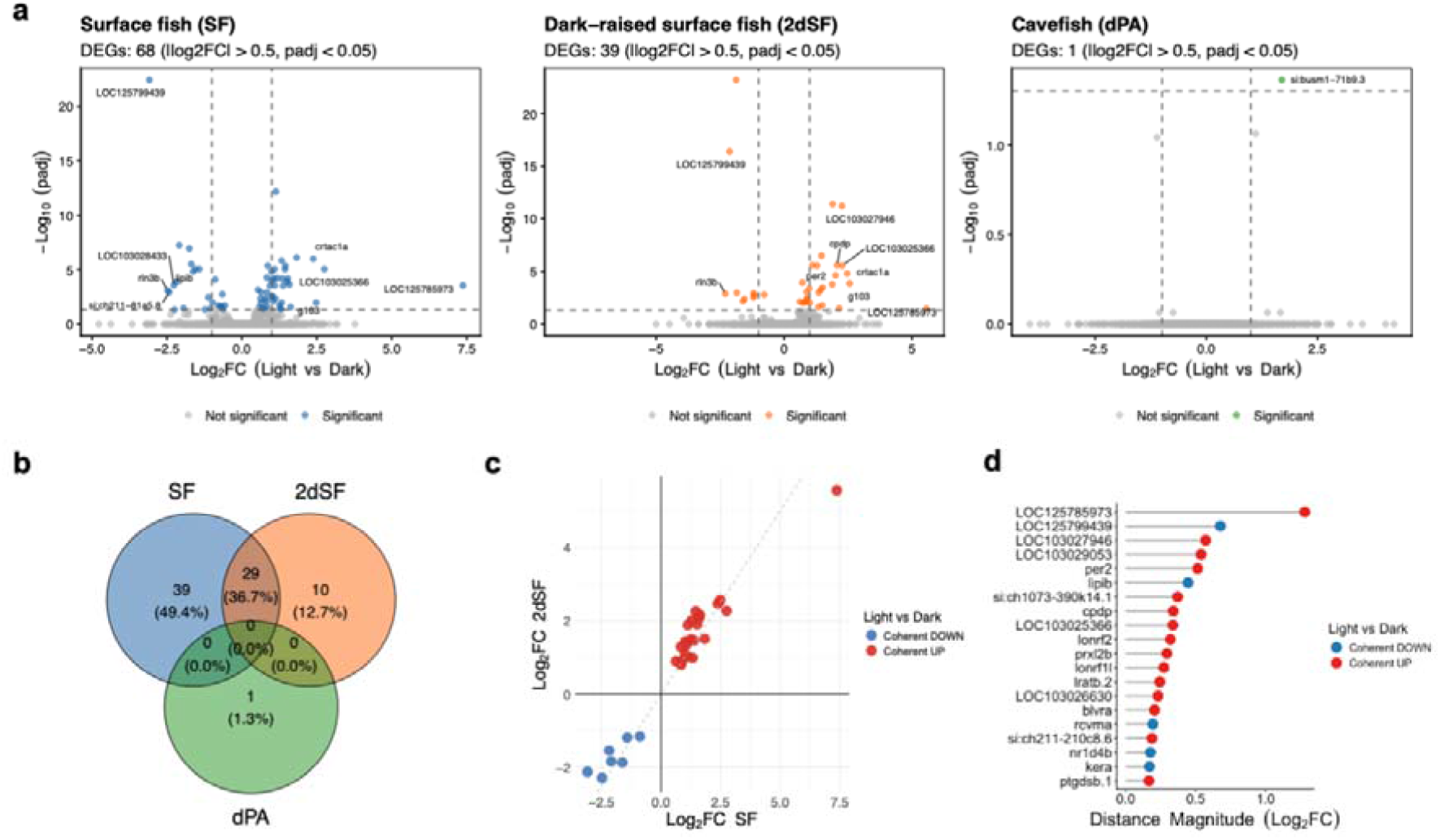
Light-induced transcriptomic plasticity across *A. mexicanus* SF, 2dSF, and dPA. Differentially expressed genes (DEGs) were identified by comparing light vs. dark conditions within each group (|log_2_FC| > 0.5, padj < 0.05). (A) Volcano plots displaying (DEGs) in response to light in SF, 2dSF, dPA. (B) Venn diagram illustrating the overlap of environment-responsive DEGs between the three groups. (C) Correlation plot of log2 fold changes between SF and 2dSF for shared light-responsive genes. (D) Lollipop plot ranking the top SF and 2dSF shared genes by their response magnitude (log2 fold change). Colors indicate the direction of the response, categorized as “Coherent UP” (red) or “Coherent DOWN” (blue) based on their consistent behavior across populations.

In contrast, 2dSF showed substantially attenuated responsiveness, with only 39 DEGs (Fig 5A, middle panel). More than half of SF-responsive genes (57%) were no longer responsive in 2dSF (S3 Table), indicating rapid loss of environmental sensitivity. Most 2dSF-responsive genes (74%) overlapped with SF-responsive genes (Fig 5B) (S4 Table) and exhibited consistent direction (Fig 5C) and effect sizes (Fig 5D). These 29 shared genes included circadian and photic response components (*per2, cry1a, crtac1a, gt03*), indicating that the core plastic response in light-signaling pathways is preserved in direction and intensity even after multigenerational darkness exposure. A small subset of 2dSF-specific DEGs (10) (S5 Table) was associated with cellular protection and mitochondrial function, potentially reflecting compensatory responses or novel regulatory connections emerging under altered developmental conditions.

In contrast, Pachón cavefish showed an almost complete absence of environment-induced transcriptional responses, with a single lineage-specific DEG (Fig 5A, right panel). This gene, *si:busm1-71b9*.*3*, is predicted to enable beta-catenin and cadherin binding activities and does not represent conservation of ancestral plasticity pathways. Overall, the reduction from 68 DEGs in SF to a single responsive gene in PA indicates that cavefish have almost entirely lost light-induced transcriptional responsiveness.

### Stable transcriptional programs associated with sustained darkness exposure

To characterize stable transcriptional divergence between SF and PA, and to determine how sustained darkness modifies transcriptional signature in 2dSF, we analyzed global gene expression across all groups on a subset of genes that did not respond to light/dark switches (|log_2_ fold change| < 1 and adjusted *P* > 0.05 for LL vs. LD and DD vs. DL comparisons). This approach identified group-specific transcriptional signatures that persist independently of short-term environmental effects.

This analysis identified 14,658 DEGs, revealing distinct expression modules across SF, 2dSF and dPA (Fig 6A). Eight major expression clusters were identified and categorized into maladaptive, adaptive and PA-specific patterns (Fig 6B). Three clusters (2, 3, and 6) displayed a putatively maladaptive pattern, characterized by exaggerated transcriptional responses in 2dSF that were not maintained in PA. Clusters 2 (2554 genes) and 3 (720 genes) were downregulated in PA, but upregulated in 2dSF relative to SF (S6 and S7 Tables, respectively). They were significantly enriched in immune genes and, unexpectedly, in transcripts related to visual perception, including several key factors such as *six6a, pax6a*, and *rx3* (Fig 6C). Cluster 6 (411 genes) encompassed downregulated genes in 2dSF, with functions in detoxification and lipid metabolism (Fig 6C; S8 Table).

**Figure 6.**
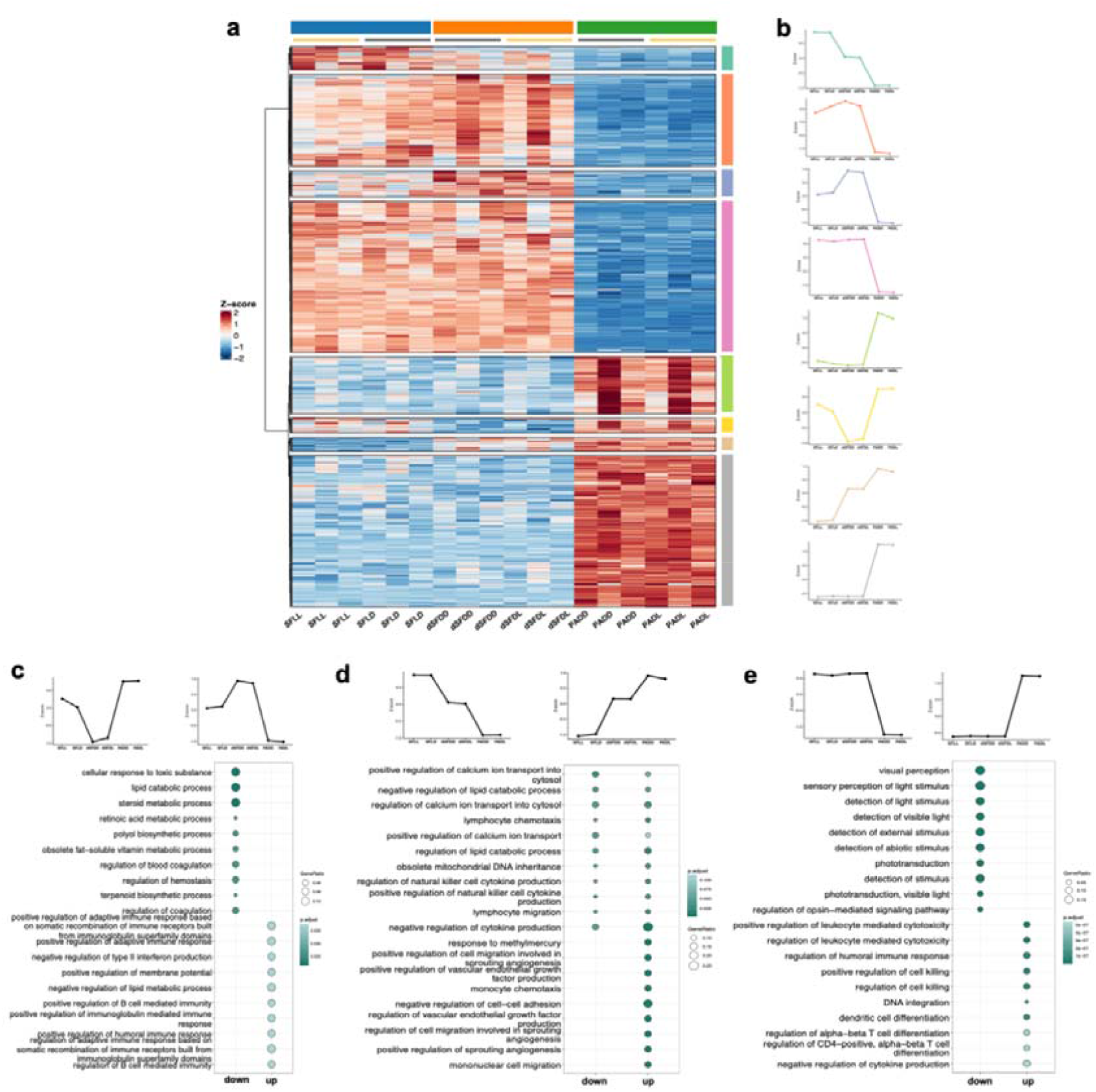
Transcriptomic divergence across *A. mexicanus* SF, 2dSF, and dPA. Analysis includes genes differentially expressed among groups (|log_2_FC| > 1, padj < 0.05) but not responsive to short-term light conditions. (A) Heatmap showing standardized expression levels (Z-scores) for genes differentially expressed among groups (SF, 2dSF, and dPA) but not responsive to short-term light conditions. Rows represent individual genes, organized by hierarchical clustering to identify modules with similar group-specific profiles. Columns represent experimental groups: surface fish (SF; SFLL, SFLD; blue), dark-raised surface fish (2dSF; 2dSFDD, 2dSFDL, orange), and Pachón cavefish (dPA; dPADD, dPADL, green). Color scale indicates relative expression (red, high; blue, low). (B) Mean expression profiles for each cluster. (C–E) Functional enrichment of biological processes for clusters showing (C) maladaptive patterns, (D) adaptive patterns, and (E) PA-specific expression patterns. In dot plots (C-E), the x-axis indicates direction of change, circle color represents adjusted P-values, and circle size indicates GeneRatio (proportion of genes in the cluster belonging to a specific GO category).

Two adaptive clusters (1 and 7; with 632 and 372 genes, respectively) showed stepwise expression shifts from SF, through 2dSF to PA consistent with the evolutionary transition from surface to cave environments. Cluster 1 (632 genes) exhibited a downregulation trend (SF > 2dSF > dPA) and was enriched for genes involved in ion transport and lipid metabolism (S9 Table) (Fig 6D). Conversely, cluster 7 (372 genes) showed a progressive upregulation (SF < 2dSF < PA) linked to regulation of cell migration and heme oxidation (S10 Table) (Fig 6D). Notably, the adaptive shifts associated with prolonged light deprivation suggest that some aspects of cavefish transcriptional programs including lipid metabolism and heme and ion homeostasis emerge in ancestral surface fish after only two generations of exposure to darkness.

Finally, cave-specific patterns were characterized by strong divergence in PA relative to both SF and 2dSF (SF≈2dSF≠PA). Cluster 4 (4199 genes) exhibited a pronounced downregulation in PA and was predominantly associated with visual and phototransduction pathways (S11 Table) (Fig 6E). In contrast, clusters 5 (1566 genes) and 8 (4204 genes) showed cave-specific upregulation, and were enriched in immune-related functions, (S12 and S13 Tables) (Fig 6E). These clusters indicate cavefish-specific specializations, including the loss of visual system and the evolution of specific immune response pathways.

## DISCUSSION

The suite of plastic responses documented here and by Bilandžija et al. (2020) indicates that the developmental programs underlying specialized phenotypes of *A. mexicanus* cave fish (CF) were already present, though environmentally regulated, in the ancestral surface population (SF). The central result of this study is that plasticity changes over remarkably short timescales. Both trait expression and patterns of environmental responsiveness are modified in surface fish after only two generations of sustained exposure (2dSF). This is consistent with phenotypic accommodation (1,11,18) which posits that environmentally induced phenotypes are modified and refined across generations. Empirical demonstrations of such rapid accommodation remain rare in natural systems (19,20), particularly under ecologically relevant conditions (6). In our study darkness was used as it is a defining and persistent feature of all subterranean habitats. Importantly, plastic responses are reshaped but nevertheless remain distinct from evolved cavefish values. Therefore, plasticity can generate rapid phenotypic change, but accommodation across generations reshapes how organisms ultimately respond to the environment. In contrast, cavefish phenotypes remain largely insensitive to environmental conditions, consistent with genetic assimilation. Together, these results support a model in which environmental cues generate plastic changes, accommodation reshapes developmental responses across generations, and canalized phenotypes represent an evolved endpoint.

Our results show that plasticity is heterogeneous. Across both morphological and molecular levels, the majority of responses in SF deviated from cavefish values, consistent with maladaptive plasticity being a pervasive biological response (21,22). Interestingly, classical cave-associated traits such as eyes and pigmentation, known to regress across diverse cave-adapted taxa, also showed maladaptive responses, corroborating our previous findings in adult *Astyanax* SF (23) and in the cave population of cyprinid *Telestes karsticus* (24). We therefore hypothesize that negative selection on maladaptive plasticity in eyes and pigmentation may not only modify plastic responses but also drive evolutionary reduction or loss of the affected traits. Maladaptive plasticity has been proposed to evolve most rapidly, as strong negative selection is expected to act on phenotypes that deviate from the local optimum (19,25–27). This scenario is consistent with eye regression in cavefish, one of the most dramatic morphological transformations documented in vertebrates over relatively short evolutionary timescales (28).

Many other phenotypes showed putatively maladaptive plasticity. Body and head height, and variables related to jaws and snout (JAHS and SNM) shifted in the appropriate direction, but overshot cavefish values. Of all the morphometric variables, only head length showed the direction and magnitude of plasticity optimally aligned with the cavefish phenotype. At the molecular level, approximately 1,000 genes were associated with adaptive changes compared to around 3,600 genes associated with maladaptive responses. Although direct comparison of gene numbers must be interpreted cautiously, because genes do not act in isolation but operate within tightly regulated, modular genetic networks (29), our results are consistent with strong maladaptive responses reported in other systems (19,26,27). The molecular basis of maladaptive plasticity is particularly evident in visual system genes, where 2dSF showed paradoxical upregulation of transcription factors associated with eye development (*six6a, pax6a, rx3*). Possibly the developmental programs evolved under surface conditions attempt to rescue visual function when normal photoreceptor signaling is absent (30). This molecular overcompensation may contribute to the maladaptive increase in eye diameter observed in 2dSF and in retinal layers’ thickness in adult *A. mexicanus* and *T. karsticus* (23,24). In PA these transcription factors are silenced, illustrating the progression from molecular overcompensation and morphological overshoot to transcription silencing and trait loss during the refinement of initially maladaptive responses into stable adaptive states.

Although adaptive responses were less prevalent in our study, anesthesia resistance represents a striking example of adaptive plasticity that undergoes rapid accommodation. The finding that 2dSF achieve the level of resistance comparable to naturally evolved cavefish within only two generations suggests that the neural circuitry underlying altered consciousness and arousal states is readily modified by prolonged darkness. Given the extensive nervous system reorganization documented in cavefish, including sleep reduction, altered neurotransmitter profiles, and enhanced sensory processing (31,32), the rapid accommodation of anesthesia resistance may reflect broader neuroplastic capacity that facilitated cave colonization in *A. mexicanus*.

Gene expression clusters showing adaptive accommodation across SF, 2dSF and PA include key metabolic regulators, suggesting that darkness alone, in the absence of nutrient deprivation or other cave-associated selective pressures, can trigger coordinated metabolic reprogramming. This is consistent with our previous finding that darkness induces starvation resistance in SF (23). Such responses may represent anticipatory plasticity (33), whereby an environmental cue (darkness) triggers physiological preparation for reliably associated, but not yet encountered conditions (12), such as nutrient limitation resulting from the absence of primary production. Natural selection would favor genotypes in which darkness functions as a predictive cue for metabolic state. This interpretation is supported by evidence that light regulates metabolism independently of feeding status in teleost fishes. Photic signals directly modulate metabolic gene expression through circadian and neuroendocrine pathways (34,35), enabling darkness to activate starvation-preparatory programs even under well-fed laboratory conditions.

Several morphometric traits and numerous genes showed no induction in 2dSF despite being distinct in cavefish (e.g. body width and otic traits), indicating phenotypes that are not environmentally inducible under this cue or developmental stage. At the molecular level more than two thirds of genes were differentially regulated in PA compared to both SF and 2dSF. The cave-specific transcriptional signature reveals expected downregulation of phototransduction pathways but also a broader reorganization of immune function, consistent with previous reports of a shift from innate to adaptive responses in cavefish (36).

Surface fish phenotypes were structured by long- and short-term environmental history and exhibited widespread plasticity, whereas cavefish formed a largely invariant cluster regardless of rearing conditions. Only a subset of traits showed detectable environmental responsiveness in PA, in stark contrast to the widespread plasticity observed in SF, and those traits exhibited markedly fewer among-group differences. No instances were observed in which a trait was non-plastic in SF but plastic in PA. Conversely, all traits that retained measurable plasticity in PA were also plastic in SF. Therefore, cavefish evolution did not involve the emergence of novel plastic traits, but rather modification or reduction of ancestral plastic responses.

Notably, all morphometric traits that lost plasticity in PA exhibited maladaptive plasticity in SF. These included eye traits and pigmentation, which increased in darkness, whereas body and head height showed plastic shifts in the appropriate direction but overshot PA values. This pattern is consistent with maladaptive ancestral plasticity being selectively reduced during cave evolution. However, not all traits showing maladaptive responses in SF became canalized in PA. Craniofacial traits (JAHS and SNM), for example, retained partial plasticity in cavefish, possibly reflecting ongoing functional demands. In cave environments, food availability is limited and unpredictable, potentially favoring retention of developmental flexibility in feeding apparatus.

These results align with models of adaptation in which rapid plastic responses initially move phenotypes toward a new optimum, followed by slower genetic assimilation and reduction of plasticity (11). Under this framework, traits exhibiting imprecise or exaggerated plasticity may be preferentially targeted for canalization, whereas traits with functionally advantageous flexibility may retain responsiveness. Together with previous empirical studies, our results suggest that maladaptive plasticity might be preferentially and rapidly targeted during the evolution of novel phenotypes (19,25–27).

The global pattern of progressive plasticity loss was also evident at the transcriptomic level. SF displayed the most extensive transcriptional plasticity, PA almost none, and 2dSF were intermediate. Most light-responsive genes in SF but environmentally unresponsive in PA, were preserved in 2dSF with consistent directionality, including core circadian regulators (*per2, cry1a*), indicating persistence of central regulatory architecture (37–39). In contrast, genes linking photic input to downstream physiological processes lost responsiveness early, including components of retinal melatonin signaling (*aanat1*) and visual chromophores regeneration in response to photobleaching (*rpe65a, rlbp1b, rdh5*), consistent with early activation of molecular programs associated with eye regression. Similarly, complement component C3, which mediates opsonization of photoreceptor outer segments and accelerates apoptotic cell clearance in retinal degeneration models (40), showed early loss of environmental responsiveness in 2dSF and constitutive upregulation in cavefish, potentially reflecting immune remodeling in the absence of photoreceptor outer segments (41). Together, these patterns indicate rapid erosion of environmentally responsive gene regulation alongside stabilization of cave-associated expression programs, in some cases within only a few generations.

Several limitations of this study warrant consideration. Plasticity was assessed at the larval stage, where shifts in trait means may be more detectable than changes in reaction norm slopes, as many traits are still undergoing rapid growth and differentiation. Compared to our previous work in adults, larvae showed a higher frequency of maladaptive responses, possibly reflecting early developmental trajectories that may be refined later in ontogeny or across generations. However, the most pronounced patterns of maladaptive plasticity, particularly in eyes and pigmentation, are consistent between larvae and adults (23), supporting the robustness of these findings.

A second consideration relates to differences in experimental design between morphotypes. Surface fish were phenotyped after two generations of dark rearing, whereas cavefish were phenotyped after a single generation. This reflects both the longer generation time of cavefish and prior evidence that cavefish phenotypes are largely unresponsive to environmental variation (23). Given this established environmental robustness, extending cavefish rearing across additional generations was unlikely to yield qualitatively different outcomes, while substantially delaying the study. Our design therefore prioritized detecting rapid transgenerational changes in the plastic surface form, while using cavefish as a reference for a canalized phenotypic regime.

Finally, we manipulated a single environmental factor, darkness, whereas natural cave environments combine multiple ecological challenges, including limited food, hypoxia, and altered biotic interactions. Consequently, the plastic and multigenerational responses observed here likely represent only a subset of the changes that occur during cave colonization. Importantly, detectable shifts in both trait expression and plasticity emerged despite the simplicity of the manipulation and the absence of strong selection in our laboratory setting, where fish were reared under benign conditions with ample food and optimal water parameters. This suggests that the observed rapid phenotypic accommodation under sustained environmental change in the laboratory may be even further accelerated when multiple ecological factors act in concert in caves. Consistent with this, hypoxia alone induces changes in eye size, erythropoiesis, heart asymmetry, and other traits associated with Sonic hedgehog signaling (42,43), highlighting the potential for interacting environmental cues to amplify and integrate plastic responses during cave adaptation (44).

Ancestral plasticity induced by darkness and other cave-related environmental cues likely played a pivotal role in the initial survival of *Astyanax* populations that colonized caves. Our evolutionary experiment, in which generations of ancestral surface-dwelling *Astyanax* are raised in complete darkness, is uniquely suited to track phenotypic responses during the earliest stages of cave-like environmental exposure. Plasticity appears exploratory rather than optimized: it biases phenotypes in a particular direction but lacks precision and frequently deviates from the evolved optimum. Such heterogeneous responses do not diminish the evolutionary importance of plasticity; instead, they generate substantial phenotypic variation which provides substrate for refinement through accommodation and selection acting on both plastic outcomes and underlying traits. The rapid phenotypic accommodation documented here provides a potential mechanistic explanation for the remarkably fast evolution observed in *Astyanax* cavefish (30–200 thousand generations; (13,14).

By empirically separating immediate plasticity, rapid accommodation, and canalized phenotypes within a single system, *Astyanax* offers a framework for understanding how plastic responses can bias phenotypic trajectories during evolutionary change. This framework may help explain adaptation to extreme environments across different taxa and their convergent evolution in response to similar cave selection pressures worldwide. In this sense, plasticity may be not just a transient response, but also a facilitator of evolutionary innovation.

## MATERIALS AND METHODS

### Fish husbandry and experimental setup

*Astyanax mexicanus* surface fish (Texas population; SF) and cavefish (Pachón population; PA) (Fig 1A) were originally obtained from the Jeffery Laboratory (University of Maryland). All experimental protocols were approved by the Institutional and National Bioethics Committees (KLASA: UP/I-322-01/22-01/42; URBROJ 525-09/589-23-4) and adhered to international and national ethical guidelines for animal research.

Fish were raised under standardized conditions with 10% daily water changes and regular monitoring of water quality: temperature was maintained at 22 ± 1°C, pH at 7.0–7.4, water conductivity at 700-800 uS. Two opposing light conditions were used: a standard 12:12-hour light:dark photoperiod (L), and complete darkness (D; fish handled under dim red light when necessary). All other environmental conditions were kept identical across treatments. Fish in L and D were housed in separate recirculating systems. Artemia feeding began at 7 days post-fertilization (dpf) for experiments involving fish older than this stage.

To explore the role of plasticity and genetic accommodation in evolving populations, we allowed SF to breed freely under conditions of complete darkness for one generation. Generation zero was derived from SF and PA embryos transferred to D within 24 hours post-fertilization (hpf). Their offspring (dSF and dPA) was never exposed to light. For this study we used four experimental groups: SF in L – surface fish maintained under standard light/dark conditions; 2dSF – the second generation of dark-raised surface fish, derived from dSF parents, PA in L and dPA - the first generation of dark-raised Pachón cavefish in D. Experimental families were generated by in vitro fertilization (IVF). IVF ensured precise control over genetic background and allowed us to disentangle the environmental from genetic effects. Gametes were collected by gently pressing the abdomens of males and females from SF in L, dSF in D, and PA in L or D. Eggs and sperm were mixed in Petri dishes with system water. Fertilized embryos were then randomly assigned to either L or D conditions within 24 hpf (Fig 1B) resulting in a full factorial experimental design. In total 8 groups were used for phenotyping, offspring raised in the same conditions as their parents SFLL, 2dSFDD, PALL and dPADD and their siblings raised in alternative environmental conditions SFLD, 2dSFDL, PALD and dPADL.

Specialized cave traits involve rearrangements across levels of organization, so we assayed a range of phenotypes including gross morphology, pigmentation, nervous system and gene expression. We were interested in both the changes of phenotypes between our respective groups and the changes in plasticity across generations of our experimental fish. All fish older than 7 dpf used in experiments were fasted for 24 hours prior to testing to standardize metabolic state. No larvae were excluded from analysis unless they exhibited gross developmental abnormalities.

### Phenotyping

#### Morphometry

At 7 dpf approximately 30 larvae from each experimental group were anesthetized using 0.1 g/L tricaine methanesulfonate (MS-222; Sigma-Aldrich, cat. #A5040, USA) and imaged using a Canon digital camera EOS 250D mounted on a dissecting microscope. The camera settings, lighting, and magnification were optimized during preliminary trials and kept constant across all groups to ensure image comparability. Each larva was photographed from the dorsal and left lateral view. Morphometric measurements were performed using ImageJ software (45). All measurements were conducted by the same researcher using pre-defined anatomical landmarks to ensure consistency in trait measurements in both SF and PA groups (S1 and S2 Figs).

Twenty-two original traits were measured from each individual: standard length (SL), anteroposterior eye diameter (APED), dorsoventral eye diameter (DVED), snout length (SNL), snout length from the center of the eye (SNLC), head height (HH), body height (BH), sagitta otolith diameter (EARSOD), distance between lapillus and sagitta otoliths (EARLS), and ear diameter (EARD) were measured from the lateral photographs. Standard length (SL), left eye width (LEW), left eye oblique diameter (LED), right eye width (REW), right eye oblique diameter (RED), jaw size (JS), distance between the eyes (DEYE), head width dorsal (HW), body width dorsal (BW), head length (HL), left pectoral fin length (LFL), and right pectoral fin length (RFL) were measured from dorsal photographs.

To remove the effect of body size on morphometric variables prior to statistical analysis, an allometric transformation was applied. For each morphometric variable, the allometric growth coefficient (b) was estmated by regressing the natural logarithm of the measurement against the natural logarithm of standard length (SL) using ordinary least-squares linear regression. Each measurement was then adjusted to the overall mean standard length 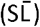 of the pooled sample using the formula: 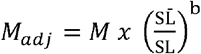, where M_adj_ is the size-adjusted measurement, and M is the original measurement (46–48). By adjusting all measurements to a common body size, this transformation ensures that observed differences among groups reflect variation in body shape rather than allometric scaling effects. The size-adjusted measurements were subsequently used in all downstream statistical analyses.

Canonical discriminant function analysis (DFA) was applied to examine morphometric differentiation among experimental groups. Given the high correlation among certain variables, composite variables were created to reduce multicollinearity. Composite variables were generated via dimensionality reduction (PCA): SNM (snout measurement) was derived from SNL and SNLC, BHH (body and head height) from HH and BH, JAHS (jaw and head size) from JS and HW, and the composite variable FINS from LFL and RFL. Twelve variables were included in the final DFA: eight individual (APED, DVED, EARSOD, EARLS, EARD, DEYE, BW, HL) and four composites (SNM, BHH, JAHS, FINS).

DFA analyses were performed at multiple levels. First, a global DFA including all eight groups (four rearing treatments × two populations; N=239) assessed overall multivariate structuring. A complementary DFA was conducted separately for cavefish (PA, N=119) and surface fish (SF, N=120) to understand the responses of each morphotype. Finally, a specific analysis compared the three groups from their natural or long-term rearing conditions, to quantify phenotypic shifts in 2nd generation of dark-reared SF (2dSFDD) relative to SF reared under standard conditions (SFLL) and evaluate their proximity to the evolved cave phenotype (dPADD). The simultaneous entry method was used for all variables. Statistical significance of discriminant functions was tested using Wilks’ lambda and chi-square test. To test the robustness of DFA and ensure the lack of overfitting, we assessed the classification accuracy using leave-one-out cross-validation.

#### Pigmentation

The same SF larvae used for morphometric measurements were photographed on 2x magnification to quantify pigmentation. Quantification was done using ImageJ. Pigmentation in SF was quantified by calculating the percentage of the polygonal region covered by melanophores. Images were converted to 8-bit format, and the polygonal region was manually drawn to include the body area posterior to the yolk sac and swim bladder (S3 Fig). An ImageJ threshold tool was applied to mark the melanophores enabling calculation of the percentage area. Minimum threshold value was set to 0, while universal maximum threshold value was used on all images. Universal maximum threshold was calculated as a mean of manually selected maximum thresholds on the random subset of images. Polygonal regions were stored for each image, enabling re-analysis. As PA are albino, L-DOPA pigmentation rescue assay was done to quantify the melanoblasts (49–51). Seven dpf PA larvae were fixed in 5% formalin in phosphate-buffered saline (PBS) for one hour at room temperature. Following fixation, larvae were washed 5 times for 5 minutes in PBS, and then stained using 0.1% L-DOPA solution (Sigma-Aldrich, USA, cat # D9628) in phosphate buffer for 6 hours at 30°C to visualize melanoblasts. After staining, fish were rinsed in PBS and imaged under standardized conditions. The number of pigment cells was manually counted using the ImageJ Cell Counter plugin along the lateral surface of each fish, posteriorly to the yolk sac and swim bladder to the tip of the tail fin (S3 Fig). Different protocols for PA and SF were used as rescued melanoblasts in PA are too small for detection with ImageJ threshold tool, while melanophores in SF are too dense for manual counting.

The Shapiro-Wilk test confirmed that data distributions were normal in both the SF and PA datasets (p > 0.05). Levene’s test showed significant heterogeneity of variances for the percentage of pigmented area in SF (F = 8.34, p < 0.001), so Welch’s ANOVA was used, followed by Games–Howell post-hoc comparisons. In contrast, the assumption of homogeneity of variances was met for the number of melanoblasts in PA (F = 0.428, p = 0.734), so a one-way ANOVA was used to compare groups.

#### Anesthesia resistance

We measured anesthesia resistance as a proxy for the level of nervous system activity (52). Individual larvae were acclimated in 2⍰mL of fish water within well plates for one hour prior to the start of the assay. To initiate the assay, an equal volume of double-strength MS222 solution was added to each well, resulting in a final concentration of 0.1⍰g/L. Larvae were observed under a dissecting microscope until all movement ceased, including in response to gentle probing with a micropipette tip, as described previously (23).

Although the Shapiro-Wilk test confirmed normality of the data (p > 0.05), Levene’s test indicated significant heterogeneity of variances (SF: F = 8.71, p <0.001; PA: F = 3.77, p = 0.014). Therefore, Welch’s ANOVA was employed for group comparisons followed by Games-Howell post-hoc tests.

### RNA sequencing and gene expression analysis

#### RNA extraction and sequencing

Pools of 10 larvae at 7.5 dpf from each experimental group were flash frozen in liquid nitrogen and stored at –80°C until further processing. Total RNA was extracted using Qiagen AllPrep DNA/RNA/miRNA Universal Kit (Qiagen, USA, cat # 80224), following the manufacturer’s protocol. The quality and quantity of extracted RNA were assessed using gel electrophoresis, an Agilent Bioanalyzer, and fluorometry with a Denovix spectrophotometer and DeNovix RNA Quantification Kit Assay (DeNovix, USA, cat # KIT-RNA-2-NS).

500 nanograms of total RNA were used for library preparation with the Lexogen CORALL RNA-Seq V2 Library Prep Kit with poly(A) RNA selection (Lexogen, Austria, cat # 177.96), including an additional rebinding step to minimize ribosomal RNA contamination. Prepared libraries were sent to Novogene (U.K., Cambridge Sequencing Center), where library quality was assessed and sequencing was performed using an Illumina NovaSeq X Plus platform to generate paired-end reads.

#### Quality control and mapping

Quality control of the raw sequencing data was performed using FastQC (v0.12.1). Adapter sequences, low-quality bases, and short reads were removed using fastp (v0.23.4) (53), default parameters were applied. Reads containing unique molecular identifiers (UMIs) were processed using UMI-tools(v1.1.5) to extract UMI sequences and append them to the read headers (54).

Cleaned reads were aligned to the *A. mexicanus* reference genome (GCF_023375975.1) (55) using STAR (v2.7.11a) (56). Only uniquely mapped reads were retained for downstream analyses. PCR duplicates were removed based on UMI information using UMI-tools dedup. Gene-level read counts were obtained using HTSeq-count (v2.0.5) (57) in union mode, based on the corresponding gene annotation file. Finally, quality control reports from all processing steps were integrated and summarized using MultiQC (v1.19) (58) to ensure data consistency across all samples.

#### Differential expression and pattern clustering

Differential expression analysis was performed using the DESeq2 R package (v1.50.2) (59). Genes with low read counts were filtered out prior to analysis. Normalization was performed using DESeq2’s median-of-ratios method. P-values were adjusted for multiple testing using the Benjamini–Hochberg method. Analyses were performed at two levels. To quantify short-term environmental plasticity, differential expression analyses were conducted independently for each group: surface fish (SFLL vs. SFLD), second generation of dark-raised surface fish (2dSFDD vs. 2dSFDL), and dark-raised Pachón cavefish (dPADD vs. dPADL) applying the threshold |log2FC| > 0.5, p-value < 0.05. A comparison of the total number of DEGs across the three groups was visualized using ggplot2 (v4.0.1).

To isolate expression differences between groups SF vs 2dSF vs PA) from short-term plastic responses, we identified genes that were significantly differentially expressed across experimental groups (|log2FC| >1, p-value < 0.05) but showed no significant response to light within any individual group. This filtering step ensured that downstream analyses captured stable transcriptional divergence and long-term environmental effects. Filtered differentially expressed genes (DEGs), Variance Stabilizing Transformation (VST) values were Z-score normalized to standardize gene expression profiles. The expression matrix was partitioned into eight distinct clusters and plotted using ComplexHeatmap package (v2.26.0) (60). Hierarchical clustering was applied to the rows within each partition using the Ward.D2 method and Pearson correlation distances. For each of the eight identified clusters, a mean expression profile was calculated by averaging the Z-scores of biological triplicates. These gene expression trends were plotted as line graphs using ggplot2 (v4.0.1), representing the centroid expression behavior across the experimental conditions.

#### Functional Annotation

Predicted protein sequences were functionally annotated using eggNOG-mapper v.2.1.12 (61), based on the eggNOG orthology database. The analysis was performed using the emapper.py script with the protein FASTA file as input. To ensure high-quality annotations, orthology assignments were restricted to Gene Ontology (GO) terms derived from non-electronic evidence (--go_evidence non-electronic). The resulting output was subsequently parsed to facilitate downstream analysis with the clusterProfiler R package (v.4.0). Specifically, GO annotations were categorized into three main domains: Biological Process (BP), Molecular Function (MF), and Cellular Component (CC).

#### Gene Ontology (GO) Enrichment Analysis

Functional enrichment analysis was performed to identify overrepresented biological processes among differentially expressed gene sets. Analyses were conducted using the clusterProfiler v.4.0 R package (62). Statistical significance was assessed using a hypergeometric test, with p-values adjusted for multiple testing using the Benjamini-Hochberg method (adjusted p-value < 0.05). To enable comparison across gene sets, the *compareCluster* function was employed to contrast enriched terms between environment- and group-specific expression profiles.

## Acknowledgements

We are grateful to Lucia Ilijić, Iva Čupić, Nikolina Kuharić, Jana Bedek, Magdalena Grgić, Lada Jovović, and Lucija Kauf for their assistance with fish maintenance. We also thank Lucija Ćuzić, Vedrana Bučko, and Luce Pavin for their help with in vitro fertilization (IVF) and embryo cleaning. This research was funded by the Tenure Track Pilot Programme of the Croatian Science Foundation and the Ecole Polytechnique Fédérale de Lausanne (TTP-2018-07-9675; EvoDark), with funds of the Croatian-Swiss Research Programme and by the Croatian Science Foundation grant (IP-2024-05-2868; EvoEye) to H.B.

## Competing Interests Statement

The authors declare no competing interests.

## Author Contributions section

Conceptualization, design and funding: HB. Experimental work: MH, LGP, MĆ, ML, TR, HB. Statistical analyses: MĆ. Bioinformatic analyses: LGP. Original draft: HB and LGP with inputs from MĆ. All authors edited and approved the last version.

## Data Availability Statement

Raw and processed RNA-seq data from all 18 samples have been deposited in the NCBI Gene Expression Omnibus (GEO) database under accession number GSE324281 (https://www.ncbi.nlm.nih.gov/geo/query/acc.cgi?acc=GSE324281). The raw data for all other phenotypic measurements is available from the corresponding author upon reasonable request.

## Supporting Information

**S1 Figure. Morphometric traits quantified in surface-dwelling *Astyanax mexicanus* larvae**. (a) Lateral view showing standard length (SL), snout length from the center of the eye (SNLC), head height (HH), and body height (BH). (b) Dorsal view showing SL, jaw size (JS), body width (BW), and right and left pectoral fin lengths (RFL, LFL). (c) Lateral head close-up showing snout length (SNL), anteroposterior eye diameter (APED), and dorsoventral eye diameter (DVED). (d) Otic region close-up showing ear diameter (EARD), sagitta otolith diameter (EARSOD), and distance between lapillus and sagitta otoliths (EARLS). (e) Dorsal head close-up showing head length (HL), head width (HW), distance between eyes (DEYE), and right (R) and left (L) eye oblique diameters (ED) and widths (EW; RED, REW, LED, LEW). Abbreviations are used consistently throughout the supplementary figures.

**S2 Figure. Morphometric traits quantified in cave-dwelling Astyanax mexicanus larvae**. (a) Lateral view showing standard length (SL), snout length from the center of the eye (SNLC), head height (HH), and body height (BH). (b) Dorsal view showing SL, jaw size (JS), body width (BW), and right and left pectoral fin lengths (RFL, LFL). (c) Lateral head close-up showing snout length (SNL), anteroposterior eye diameter (APED), and dorsoventral eye diameter (DVED). (d) Otic region close-up showing ear diameter (EARD), sagitta otolith diameter (EARSOD), and distance between lapillus and sagitta otoliths (EARLS). (e) Dorsal head close-up showing head length (HL), head width (HW), distance between eyes (DEYE), and right (R) and left (L) eye oblique diameters (ED) and widths (EW; RED, REW, LED, LEW). Abbreviations are used consistently throughout the supplementary figures.

**S3 Figure. Pigmentation quantification in cave- and surface-dwelling *Astyanax mexicanus* larvae**.

(a) Lateral view of cavefish larvae showing manual melanophore counts performed using the ImageJ Cell Counter plugin. Counts were restricted to the lateral body surface posterior to the swim bladder, extending to the tip of the tail fin. (b) Lateral view of surface fish larvae showing the polygonal region of interest (yellow outline) used for pigmentation quantification in the same anatomical area. Melanophores were detected using the ImageJ threshold tool (red), enabling calculation of the percentage area covered.

**S4 Figure**. Structure matrix heatmap showing pooled within-group correlations between twelve discriminant variables (see S1 and S2 Figures for definitions) and four canonical functions from a combined discriminant function analysis (DFA) of all eight experimental groups. Groups included surface fish (SF), second-generation dark-reared surface fish (2dSF), Pachón cavefish (PA), and dark-reared Pachón cavefish (dPA), each reared under standard 12 h:12 h light:dark conditions or in constant darkness. Variables are ordered by absolute correlation values and grouped according to their strongest loading on each function.

**S5 Figure**. Structure matrix heatmap from a discriminant function analysis (DFA) of surface fish groups (SF and 2dSF). Pooled within-group correlations between twelve discriminant variables (see S1 and S2 Figures for definitions) and canonical functions are shown. Variables are ordered by absolute correlation values and grouped according to their strongest loading on each function. Group designations and rearing conditions are defined in S4 Figure.

**S6 Figure**. Structure matrix heatmap from a discriminant function analysis (DFA) of cavefish groups (PA and dPA). Pooled within-group correlations between twelve discriminant variables (see S1 and S2 Figures for definitions) and canonical functions are shown. Variables are ordered by absolute correlation values and grouped according to their strongest loading on each function. Group designations and rearing conditions are defined in S4 Figure.

**S1 Table**. Classification results from discriminant function analysis (DFA) of Astyanax mexicanus experimental groups. 1A cavefish groups, and 1B surface fish groups.

**S2 Table**. Classification results from discriminant function analysis (DFA) of all experimental groups.

**S3 Table**. Genes uniquely light-responsive in SF. List of the 39 genes identified as differentially expressed uniquely in surface fish (SF) when comparing SFLL and SFLD.

**S4 Table**. Conserved light-responsive genes shared between SF and 2dSF. List of the 29 genes identified as differentially expressed in response to light in both surface fish (SF) and dark-raised surface fish (2dSF).

**S5 Table**. Light-responsive in 2dSF. List of the 10 genes identified as differentially expressed uniquely in dark-raised surface fish (2dSF) when compared to 2dSFDD and 2dSFDL.

**S6 Table**. List of genes comprising cluster 2 from the group-specific transcriptomic analysis under sustained darkness exposure.

**S7 Table**. List of genes comprising cluster 3 from the group-specific transcriptomic analysis under sustained darkness exposure.

**S8 Table**. List of genes comprising cluster 6 from the group-specific transcriptomic analysis under sustained darkness exposure.

**S9 Table**. List of genes comprising cluster 1 from the group-specific transcriptomic analysis under sustained darkness exposure.

**S10 Table**. List of genes comprising cluster 7 from the group-specific transcriptomic analysis under sustained darkness exposure.

**S11 Table**. List of genes comprising cluster 4 from the group-specific transcriptomic analysis under sustained darkness exposure.

**S12 Table**. List of genes comprising cluster 5 from the group-specific transcriptomic analysis under sustained darkness exposure.

**S13 Table**. List of genes comprising cluster 8 from the group-specific transcriptomic analysis under sustained darkness exposure.

## Notes

### Competing Interest Statement

The authors have declared no competing interest.

## References

1. West-Eberhard MJ. Developmental Plasticity and Evolution. Dev Plast Evol. 2003 Mar 13;

2. Pfennig DW. Phenotypic Plasticity & Evolution. Phenotypic Plast Evol. 2021 May 31;

3. Waddington CH. GENETIC ASSIMILATION OF AN ACQUIRED CHARACTER. Evolution (N Y). 1953 Jun 1;7(2):118–26.

4. Ghalambor CK, McKay JK, Carroll SP, Reznick DN. Adaptive versus non-adaptive phenotypic plasticity and the potential for contemporary adaptation in new environments. Funct Ecol. 2007 Jun 1;21(3):394–407.

5. Ehrenreich IM, Pfennig DW. Genetic assimilation: a review of its potential proximate causes and evolutionary consequences. Ann Bot. 2016 Apr 1;117(5):769–79.

6. Levis NA, Pfennig DW. Evaluating “Plasticity-First” Evolution in Nature: Key Criteria and Empirical Approaches. Trends Ecol Evol. 2016 Jul 1;31(7):563–74.

7. West-Eberhard MJ. Developmental plasticity and the origin of species differences. Proc Natl Acad Sci U S A. 2005 May 3;102 Suppl(Suppl 1):6543–9.

8. Crispo E. The Baldwin effect and genetic assimilation: revisiting two mechanisms of evolutionary change mediated by phenotypic plasticity. Evolution. 2007 Nov;61(11):2469–79.

9. Pigliucci M, Murren CJ, Schlichting CD. Phenotypic plasticity and evolution by genetic assimilation. J Exp Biol. 2006 Jun 15;209(12):2362–7.

10. Sommer RJ. Phenotypic Plasticity: From Theory and Genetics to Current and Future Challenges. Genetics. 2020 May 1;215(1):1–13.

11. Lande R. Adaptation to an extraordinary environment by evolution of phenotypic plasticity and genetic assimilation. J Evol Biol. 2009 Jul 1;22(7):1435–46.

12. Culver DC, Pipan T. The Biology of Caves and Other Subterranean Habitats. Biol Caves Other Subterr Habitats. 2019 Apr 1;

13. Fumey J, Hinaux H, Noirot C, Thermes C, Rétaux S, Casane D. Evidence for late Pleistocene origin of Astyanax mexicanus cavefish. BMC Evol Biol. 2018 Apr 18;18(1).

14. Herman A, Brandvain Y, Weagley J, Jeffery WR, Keene AC, Kono TJY, et al. The role of gene flow in rapid and repeated evolution of cave-related traits in Mexican tetra, Astyanax mexicanus. Mol Ecol. 2018 Nov 1;27(22):4397–416.

15. Yoshizawa M. Behaviors of cavefish offer insight into developmental evolution. Mol Reprod Dev. 2015 Apr 1;82(4):268–80.

16. Kowalko J. Utilizing the blind cavefish Astyanax mexicanus to understand the genetic basis of behavioral evolution. J Exp Biol. 2020 Feb 1;223(Pt Suppl 1).

17. Bilandžija H, Abraham L, Ma L, Renner KJ, Jeffery WR. Behavioural changes controlled by catecholaminergic systems explain recurrent loss of pigmentation in cavefish. Proceedings Biol Sci. 2018 May 16;285(1878).

18. Schneider RF, Meyer A. How plasticity, genetic assimilation and cryptic genetic variation may contribute to adaptive radiations. Mol Ecol. 2017 Jan 1;26(1):330–50.

19. Ghalambor CK, Hoke KL, Ruell EW, Fischer EK, Reznick DN, Hughes KA. Non-adaptive plasticity potentiates rapid adaptive evolution of gene expression in nature. Nature. 2015 Sep 17;525(7569):372–5.

20. Levis NA, Isdaner AJ, Pfennig DW. Morphological novelty emerges from pre-existing phenotypic plasticity. Nat Ecol Evol 2018 28. 2018 Jul 9;2(8):1289–97.

21. Crispo E. Modifying effects of phenotypic plasticity on interactions among natural selection, adaptation and gene flow. J Evol Biol. 2008 Nov;21(6):1460–9.

22. Hendry AP. Chapter 11. Plasticity. Eco-evolutionary Dynamics. Princeton University Press; 2017. 276–303 p.

23. Bilandžija H, Hollifield B, Steck M, Meng G, Ng M, Koch AD, et al. Phenotypic plasticity as a mechanism of cave colonization and adaptation. Elife. 2020 Apr 1;9.

24. Čupić M, Marčić Z, Lukić M, Gračan R, Bilandžija H, Čupić M, et al. The first cavefish in the Dinaric Karst? Cave colonization made possible by phenotypic plasticity in Telestes karsticus. Zool Res 2023, Vol 44, Issue 4, Pages 821-833. 2023 Jul 18;44(4):821–33.

25. Fischer EK, Ghalambor CK, Hoke KL. Plasticity and evolution in correlated suites of traits. J Evol Biol. 2016 May 1;29(5):991–1002.

26. Ho WC, Zhang J. Evolutionary adaptations to new environments generally reverse plastic phenotypic changes. Nat Commun 2018 91. 2018 Jan 24;9(1):350–.

27. Campbell-Staton SC, Velotta JP, Winchell KM. Selection on adaptive and maladaptive gene expression plasticity during thermal adaptation to urban heat islands. Nat Commun 2021 121. 2021 Oct 26;12(1):6195–.

28. Jeffery WR. Astyanax surface and cave fish morphs. Evodevo. 2020 Jul 11;11(1):14–.

29. Van Gestel J, Weissing FJ. Is plasticity caused by single genes? Nat 2018 5557698. 2018 Mar 29;555(7698):E19–20.

30. Khoussine J, Sawant A, Gupta S, Zhai H, Shahi PK, Pattnaik BR, et al. Divergent mechanisms of neural adaptation and instability in the mammalian retina. Curr Biol. 2025 Jul 21;35(14):3381–3395.e4.

31. Soares D, Niemiller ML. Sensory Adaptations of Fishes to Subterranean Environments. Bioscience. 2013 Apr 1;63(4):274–83.

32. Jaggard JB, Lloyd E, Yuiska A, Patch A, Fily Y, Kowalko JE, et al. Cavefish brain atlases reveal functional and anatomical convergence across independently evolved populations. Sci Adv. 2020 Sep 1;6(38):3126–42.

33. Petrullo L, Morris NJ, Tharin C, Dantzer B. Harbingers of change: Towards a mechanistic understanding of anticipatory plasticity in animal systems. Funct Ecol. 2025 Nov 1;39(11):2999–3020.

34. Isorna E, de Pedro N, Valenciano AI, Alonso-Gómez ÁL, Delgado MJ. Interplay between the endocrine and circadian systems in fishes. J Endocrinol. 2017;232(3):R141–59.

35. Cowan M, Azpeleta C, López-Olmeda JF. Rhythms in the endocrine system of fish: a review. J Comp Physiol B 2017 1878. 2017 Apr 26;187(8):1057–89.

36. Peuß R, Box AC, Chen S, Wang Y, Tsuchiya D, Persons JL, et al. Adaptation to low parasite abundance affects immune investment and immunopathological responses of cavefish. Nat Ecol Evol. 2020 Oct 1;4(10):1416–30.

37. Beale A, Guibal C, Tamai TK, Klotz L, Cowen S, Peyric E, et al. Circadian rhythms in Mexican blind cavefish Astyanax mexicanus in the lab and in the field. Nat Commun 2013 41. 2013 Nov 14;4(1):2769–.

38. Mack KL, Jaggard JB, Persons JL, Roback EY, Passow CN, Stanhope BA, et al. Repeated evolution of circadian clock dysregulation in cavefish populations. PLoS Genet. 2021 Jul 12;17(7).

39. Di Rosa V, Frigato E, Negrini P, Cristiano W, López-Olmeda JF, Rétaux S, et al. Sporadic feeding regulates robust food entrainable circadian clocks in blind cavefish. iScience. 2024 Jul 19;27(7).

40. Hoh Kam J, Lenassi E, Malik TH, Pickering MC, Jeffery G. Complement component C3 plays a critical role in protecting the aging retina in a murine model of age-related macular degeneration. Am J Pathol. 2013 Aug;183(2):480–92.

41. Emam A, Yoffe M, Cardona H, Soares D. Retinal morphology in Astyanax mexicanus during eye degeneration. J Comp Neurol. 2020 Jun 15;528(9):1523–34.

42. Yamamoto Y, Stock DW, Jeffery WR. Hedgehog signalling controls eye degeneration in blind cavefish. Nature. 2004 Oct 14;431(7010):844–7.

43. van der Weele CM, Jeffery WR. Cavefish cope with environmental hypoxia by developing more erythrocytes and overexpression of hypoxia-inducible genes. Elife. 2022 Jan 1;11.

44. Ng M, Bilandžija H, Jeffery WR. Environmental hypoxia controls the evolution of cavefish heart asymmetry. bioRxiv. 2026 Jan 30;2026.01.29.702597.

45. Schneider CA, Rasband WS, Eliceiri KW. NIH Image to ImageJ: 25 years of image analysis. Nat Methods 2012 97. 2012 Jun 28;9(7):671–5.

46. Reist JD. An empirical evaluation of several univariate methods that adjust for size variation in morphometric data. Can J Zool. 2011 Jun 1;63(6):1429–39.

47. Elliott NG, Haskard K, Koslow JA. Morphometric analysis of orange roughy (Hoplostethus atlanticus) off the continental slope of southern Australia. J Fish Biol. 1995 Feb 1;46(2):202–20.

48. Lleonart J, Salat J, Torres GJ. Removing allometric effects of body size in morphological analysis. J Theor Biol. 2000 Jul 7;205(1):85–93.

49. McCauley DW, Hixon E, Jeffery WR. Evolution of pigment cell regression in the cavefish Astyanax: a late step in melanogenesis. Evol Dev. 2004 Jul 1;6(4):209–18.

50. Bilandžija H, Ma L, Parkhurst A, Jeffery WR. A Potential Benefit of Albinism in Astyanax Cavefish: Downregulation of the oca2 Gene Increases Tyrosine and Catecholamine Levels as an Alternative to Melanin Synthesis. PLoS One. 2013 Nov 25;8(11):e80823.

51. Bilandžija H, Renner KJ, Cetković H, Jeffery WR. Convergent evolution of albinism in cave animals is driven by shared mechanisms and adaptive advantages. Commun Biol 2026. 2026 Apr 15;

52. Leyden C, Brüggemann T, Debinski F, Simacek CA, Dehmelt FA, Arrenberg AB. Efficacy of Tricaine (MS-222) and Hypothermia as Anesthetic Agents for Blocking Sensorimotor Responses in Larval Zebrafish. Front Vet Sci. 2022 Mar 28;9:864573.

53. Chen S, Zhou Y, Chen Y, Gu J. fastp: an ultra-fast all-in-one FASTQ preprocessor. Bioinformatics. 2018 Sep 1;34(17):i884.

54. Smith T, Heger A, Sudbery I. UMI-tools: Modeling sequencing errors in Unique Molecular Identifiers to improve quantification accuracy. Genome Res. 2017 Mar 1;27(3):491–9.

55. Warren WC, Boggs TE, Borowsky R, Carlson BM, Ferrufino E, Gross JB, et al. A chromosome-level genome of Astyanax mexicanus surface fish for comparing population-specific genetic differences contributing to trait evolution. Nat Commun 2021 121. 2021 Mar 4;12(1):1447–.

56. Dobin A, Davis CA, Schlesinger F, Drenkow J, Zaleski C, Jha S, et al. STAR: Ultrafast universal RNA-seq aligner. Bioinformatics. 2013;29(1):15–21.

57. Anders S, Pyl PT, Huber W. HTSeq-A Python framework to work with high-throughput sequencing data. Bioinformatics. 2015;

58. Ewels P, Magnusson M, Lundin S, Käller M. MultiQC: summarize analysis results for multiple tools and samples in a single report. Bioinformatics. 2016 Oct 1;32(19):3047–8.

59. Love MI, Huber W, Anders S. Moderated estimation of fold change and dispersion for RNA-seq data with DESeq2. Genome Biol. 2014 Dec;15(12):550.

60. Gu Z, Eils R, Schlesner M. Complex heatmaps reveal patterns and correlations in multidimensional genomic data. Bioinformatics. 2016 Sep 15;32(18):2847–9.

61. Cantalapiedra CP, HerntLandez-Plaza A, Letunic I, Bork P, Huerta-Cepas J. eggNOG-mapper v2: Functional Annotation, Orthology Assignments, and Domain Prediction at the Metagenomic Scale. Mol Biol Evol. 2021 Dec 9;38(12):5825–9.

62. Wu T, Hu E, Xu S, Chen M, Guo P, Dai Z, et al. clusterProfiler 4.0: A universal enrichment tool for interpreting omics data. Innovation. 2021;2(3).

