## Supplementary figures and images for "Sustained darkness reveals rapid phenotypic accommodation and loss of plasticity in cavefish evolution"

### Supplemental Figure 1

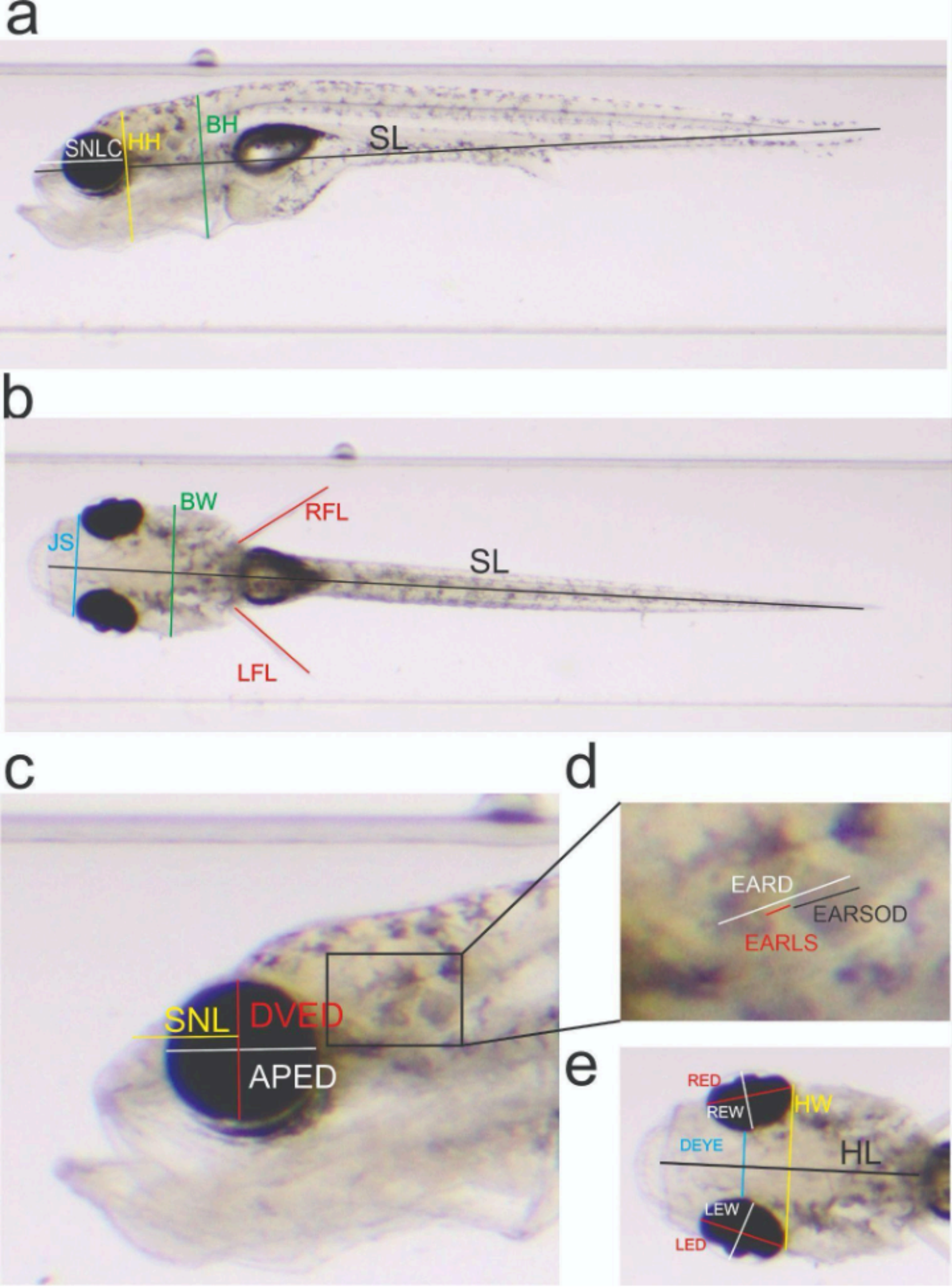

### Supplemental Figure 2

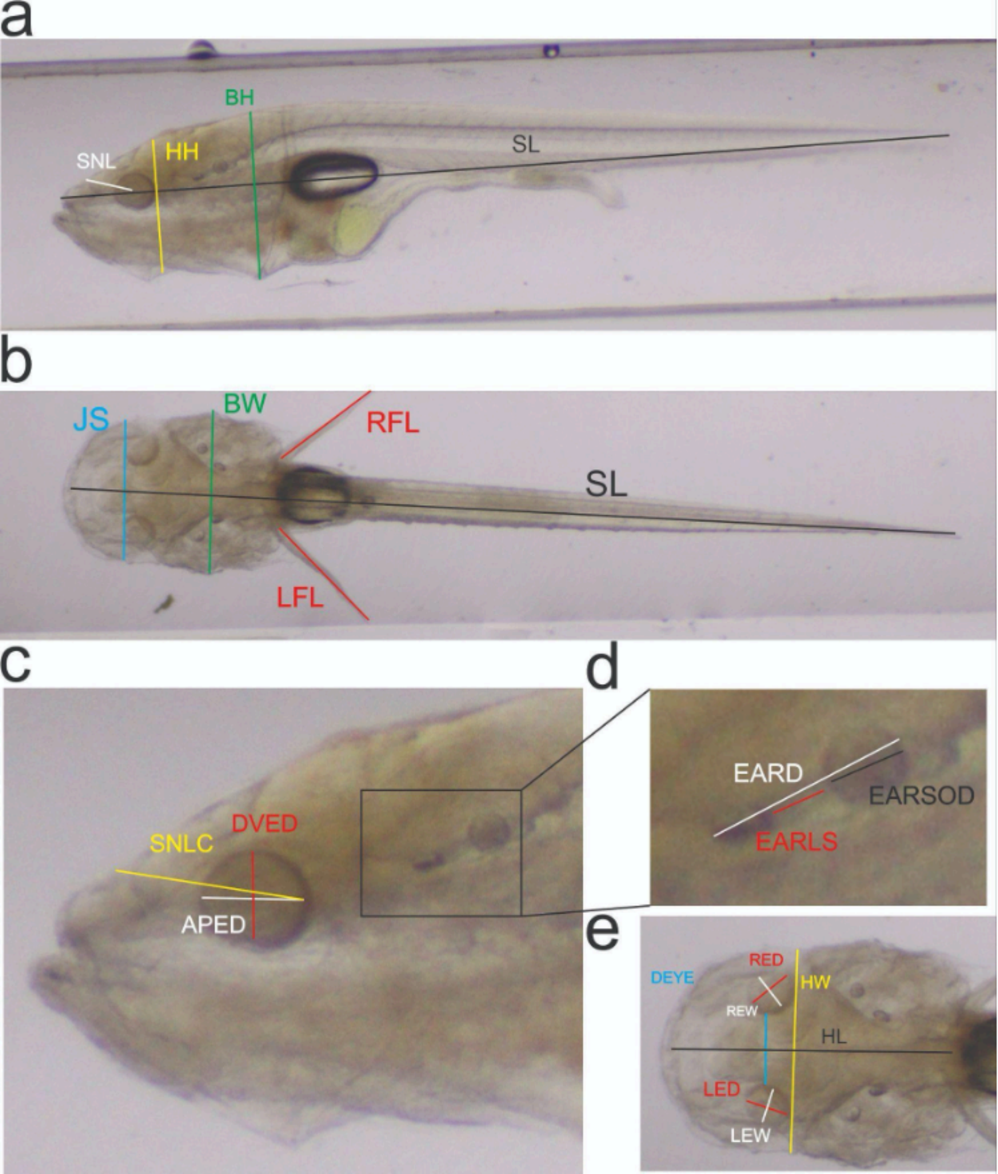

### Supplemental Figure 3

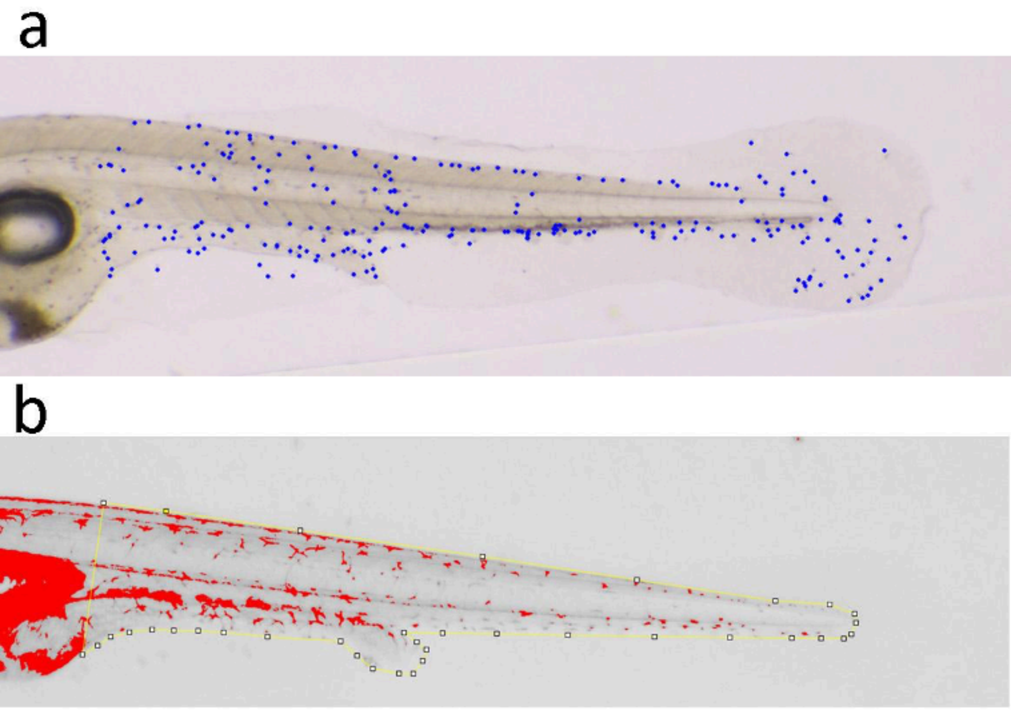

### Supplemental Figure 4

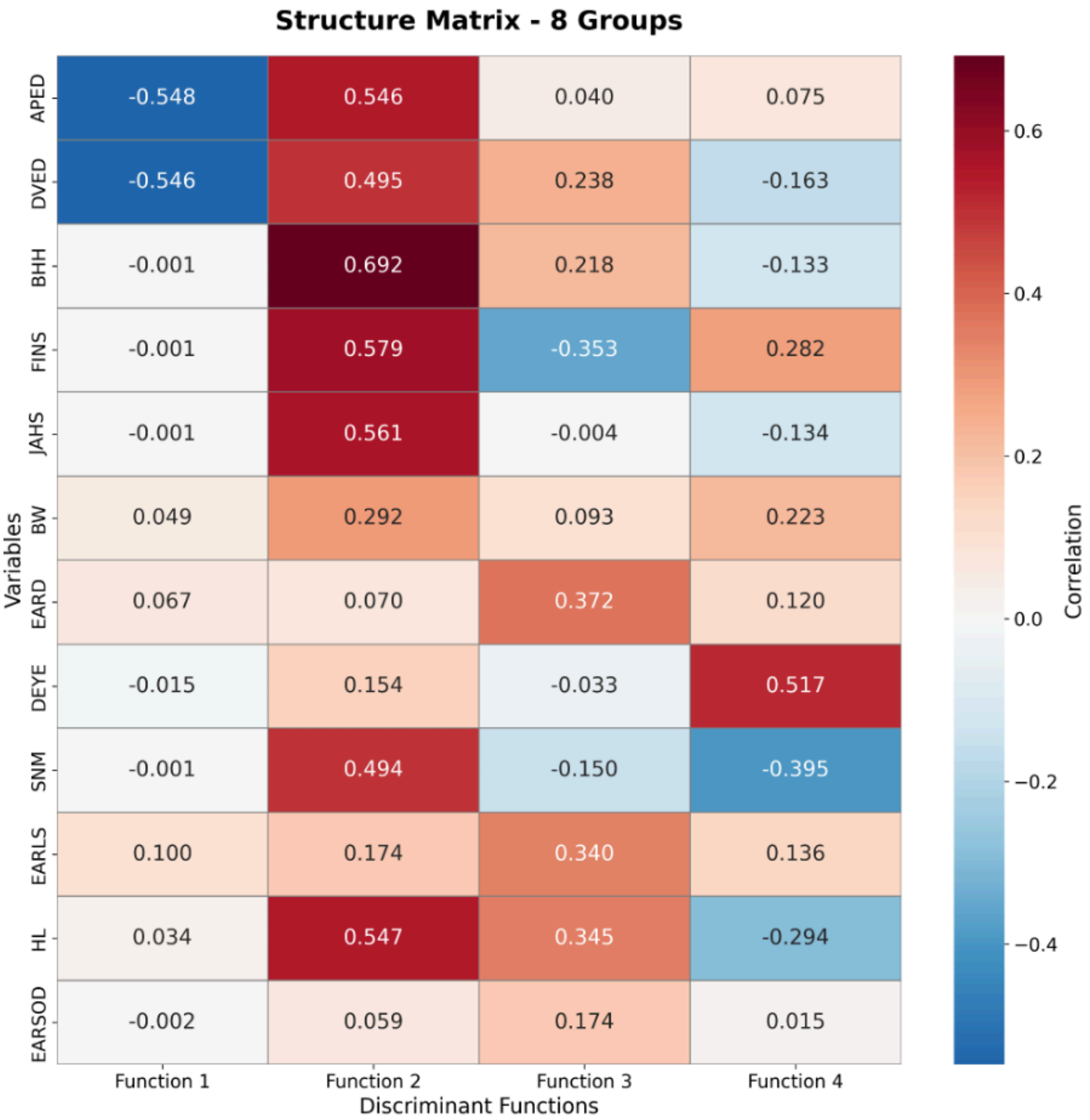

### Supplemental Figure 5

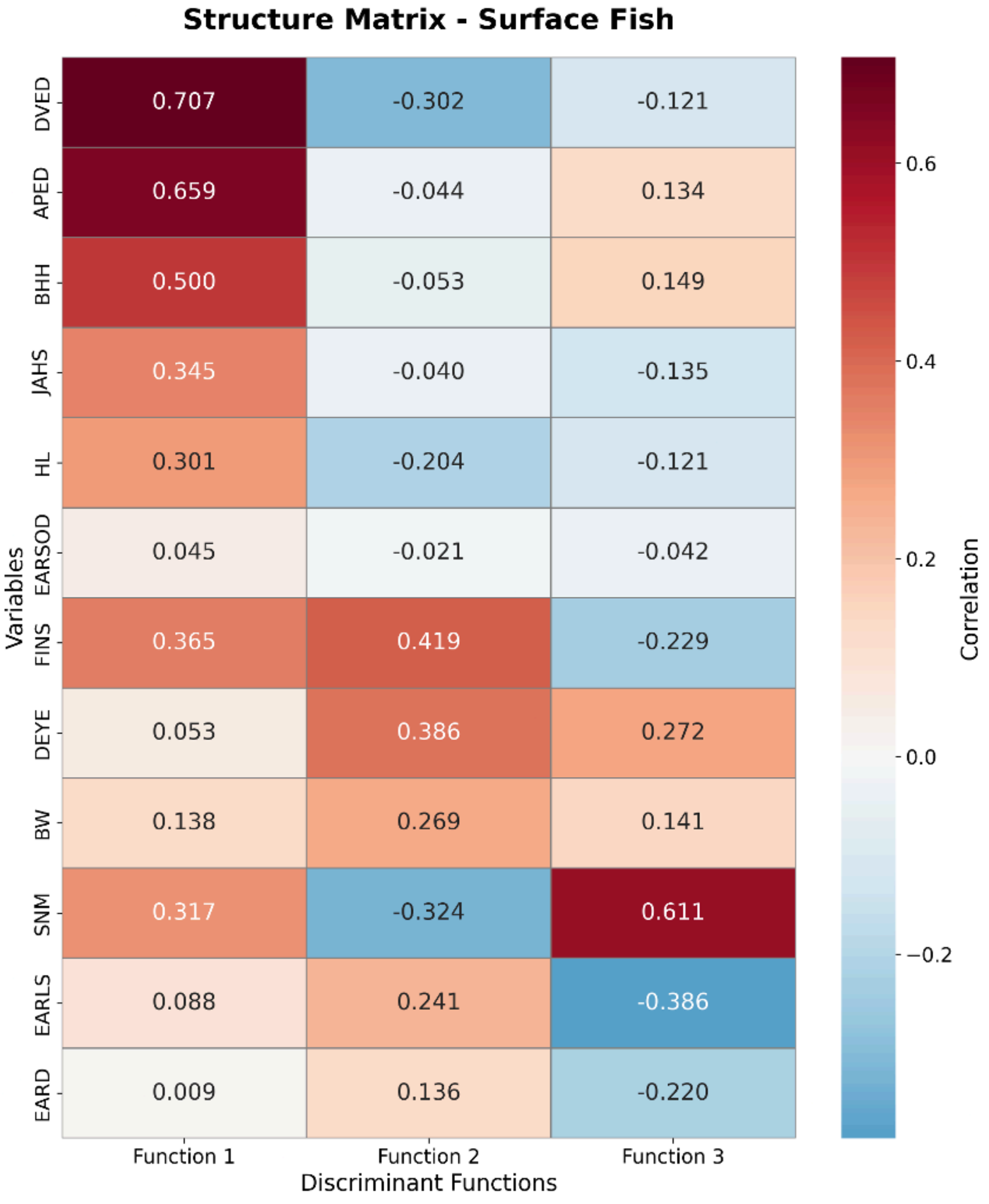

### Supplemental Figure 6

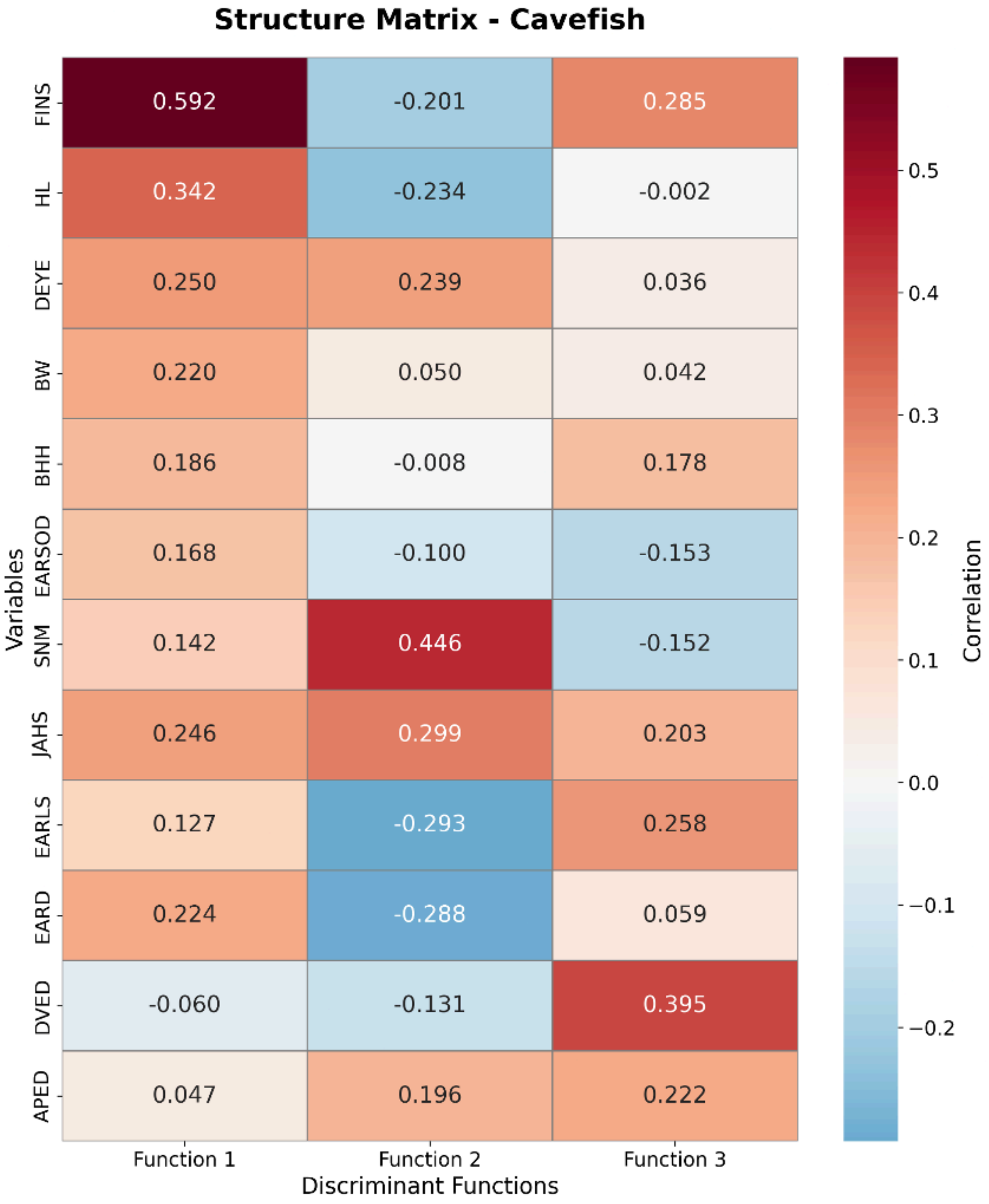
